# Mice navigate scent trails using predictive policies

**DOI:** 10.1101/2025.08.27.672631

**Authors:** Siddharth Jayakumar, Nicola Rigolli, Mohammed Mostafizur Rahman, Mackenzie Weygandt Mathis, Massimo Vergassola, Alexander Mathis, Venkatesh N. Murthy

## Abstract

Many terrestrial animals accurately follow odor trails, but their navigation strategies remain unclear. We presented mice with dynamic, non-repeating odor trails using a paper treadmill, and found that they rapidly learned to track trails effectively. Odor detections by one or both nostrils during inhalation evoked rapid, stereotyped motor responses. Blocking one nostril led to biased tracking, while disconnecting interhemispheric neural communication disrupted tracking almost entirely. We found that trail tracking is more than a reflex, with mice adapting their movements according to a memory of recently encountered trail geometry. A Bayesian model captures these behaviors, suggesting mice combine bilateral olfactory input with short-term memory to guide navigation actively. Functional perturbations and neural recordings from behaving animals implicate an early olfactory cortical region in trail tracking. These results lay the groundwork to uncover the computational architecture of sensorimotor integration in naturalistic tasks.

## Introduction

Animals use their chemical senses to interact with and explore the world. Odor-based navigation is well known in insects, birds and mammals - notably rodents and dogs - but the behavioral strategies employed remain poorly understood. By making a series of decisions using stimuli in the world and using an internal map of the environment, animals can navigate large distances. Scent trails act as navigational landmarks that enable animals to identify conspecifics and salient places like home (Draft et al., 2018; Jones and Urban, 2018; Khan et al., 2012; Porter et al., 2007; Thesen et al., 1993). However, following such trails is a nontrivial problem because they are intermittent and dynamic. Furthermore, olfaction is a proximal sense. In contrast to vision-based navigation, where distant targets and trajectory options are immediately apparent, olfactory cues provide limited information about the spatial distribution or future locations of the chemical trail.

To track scent trails on the ground or odor plumes in air, animals make characteristic zig-zag like movements, commonly called casting (Draft et al., 2018; Khan et al., 2012; Porter et al., 2007). Such casting movements are observed in flying insects and are thought to be a part of search strategies employed to discover the source (Basu and Nagel, 2024; Cardé, 2021; Reddy et al., 2022a). Moths are known to complement casting with upwind surges upon detecting odors. Such search strategies are complemented by other sensory modalities, including optomotor anemotaxis (Álvarez-Salvado et al., 2018; Demir et al., 2020; Gomez-Marin and Louis, 2014).

Surface-borne scents pose a distinct set of challenges compared to airborne odors, where dynamic olfactory landscapes arise even when plumes are continuous, and more so when they are discontinuous due to turbulent transport (Gorur-Shandilya et al., 2019; Reddy et al., 2022a). Scents adsorbed on surfaces have a more stable distribution and have far slower transient temporal fluctuations than airborne cues. However, surface-borne scents can degrade over time, may not have predictable spatial gradients, and are challenging to track when they are meandering and discontinuous. Despite these challenges, wolves, dogs and bears can track trails many miles in length with remarkable accuracy (Peters and Mech, 1975; Thesen et al., 1993; Togunov et al., 2017). To follow a scent trail, an animal must conduct an initial search to locate the scent, followed by estimation of the heading of the trail, and direct subsequent movement to track the trail. In laboratory settings, animals as diverse as rodents, ants, and humans show similarities in tracking strategies despite differences in sensorimotor constraints (Draft et al., 2018; Jones and Urban, 2018; Khan et al., 2012; Porter et al., 2007).

Strategies for odor-guided animal navigation of airborne or surface-bound cues can be distinguished based on whether they are reactive or predictive (Emonet and Vergassola, 2024; Murthy, 2024; Reddy et al., 2022a). In one formulation, animals react to immediate sensory cues or landmarks to make decisions, for example, surging upwind when encountering odors (Álvarez-Salvado et al., 2018; Demir et al., 2020; Gomez-Marin and Louis, 2014) or turning based on spatial or temporal gradients (Gardiner and Atema, 2010; Gaudry et al., 2012; Khan et al., 2012; Liu et al., 2020; Rabell et al., 2017). Many models of odor navigation rely on such reactive strategies (Belanger and Arbas, 1998; Hengenius et al., 2021; Li et al., 2001; Russell et al., 2003; van Breugel and Dickinson, 2014), which are effective when cues are frequent and reliably indicate direction. Indeed, previous studies have shown that rodents can track relatively simple odor trails in a laboratory setting, and have proposed reactive strategies involving comparisons of odor intensities across nostrils or successive samples, effectively calculating spatial or temporal gradients of odor (Khan et al., 2012; Liu et al., 2020). However, the strategies described previously rely on immediate sensorimotor responses, and do not work when odor stimuli are intermittent or absent. These reactive strategies also do not easily account for any adaptations or reliance of prior experience.

Model-based navigation strategies, in contrast, make use of prior knowledge, including recent experiences (Jayakumar and Murthy, 2022; Reddy et al., 2022b; Singh et al., 2023; Vergassola et al., 2007). These predictive strategies are particularly helpful when sensory cues are sparse, conditions under which reactive strategies may fail during long pauses in cue encounters. In particular, a recent theoretical proposal offers a predictive framework where an agent can use past encounters with the trail to estimate the trail heading direction as well as an angular sector of uncertainty about the heading (Reddy et al., 2022b). This formulation is broadly related to Bayesian inference, in which agents can balance current sensory input and priors to estimate the posterior probability of the underlying cause (Aitchison and Lengyel, 2017). Recent experiments in flies point to a role for short-term working memory of goal heading direction in odor navigation (Kathman et al., 2024; Siliciano et al., 2025). Whether and how animals, including mice use a working memory of the scent trail statistics to efficiently track uncertain trails, is unknown.

Here, we used a custom-designed paper treadmill apparatus to print long odor trails of varying geometry and combined it with quantitative behavioral analysis and modeling to show that mice combine immediate sensory information with memory as they navigate scent trails. Furthermore, we found evidence of predictive signals in neural activity of early olfactory areas.

## Results

### Mice quickly learn to follow continuous ground-based trails

Using our custom-built paper treadmill (Figure 1a), we presented mice with continuous odor trails of 2-phenylethanol, printed in different shapes and statistics (Sample videos in https://github.com/VNMurthyLab/trail_tracking/blob/main/README.md). The experimental setup uses near infrared lighting (>850nm) to enable video recording while rendering the environment largely invisible to mice. To encourage mice to follow odor trails for long periods of time, we presented chocolate flavored Ensure drops delivered randomly on the trail to increase their motivation (see Methods). Naïve mice followed odor trails even on the first day of exposure and progressively became experts at closely following trails with their snout by day 6, as evidenced by their reduced deviations (Figure 1b-e; 61.0 ± 3.8 % tracking in expert vs 7.3 ± 2.12 % in naïve, mean ± s.e.m, 13 animals, D = 0.32, p = 0.002, Kolmogorov-Smirnoff (KS) test). To select episodes of trail tracking, we defined a tracking epoch as any period during which the animal remained within 2.5 cm on either side of the trail for at least 2.5 seconds. Over days, such epochs of trail tracking increase in number and duration (Figure 1f-h; 2.7 ± 0.71 epochs in naïve vs 16.2 ± 1.29 in expert, mean ± s.e.m, 13 animals, p = 0.0002, Wilcoxon signed-rank test). The improved trail tracking metric with experience was not due to specific selection criteria (Figure S1a-c, 71.7 ± 3.03% tracking in expert vs 9.4 ± 2.59 % in naïve, mean ± s.e.m, at 3.5cm for 2.5 seconds and Figure S1d-f, 41.2 ± 5.22% in expert vs 4.9 ± 1.53% in naïve at 1.5cm for 2.5 seconds, mean ± s.e.m).

**Figure 1:**
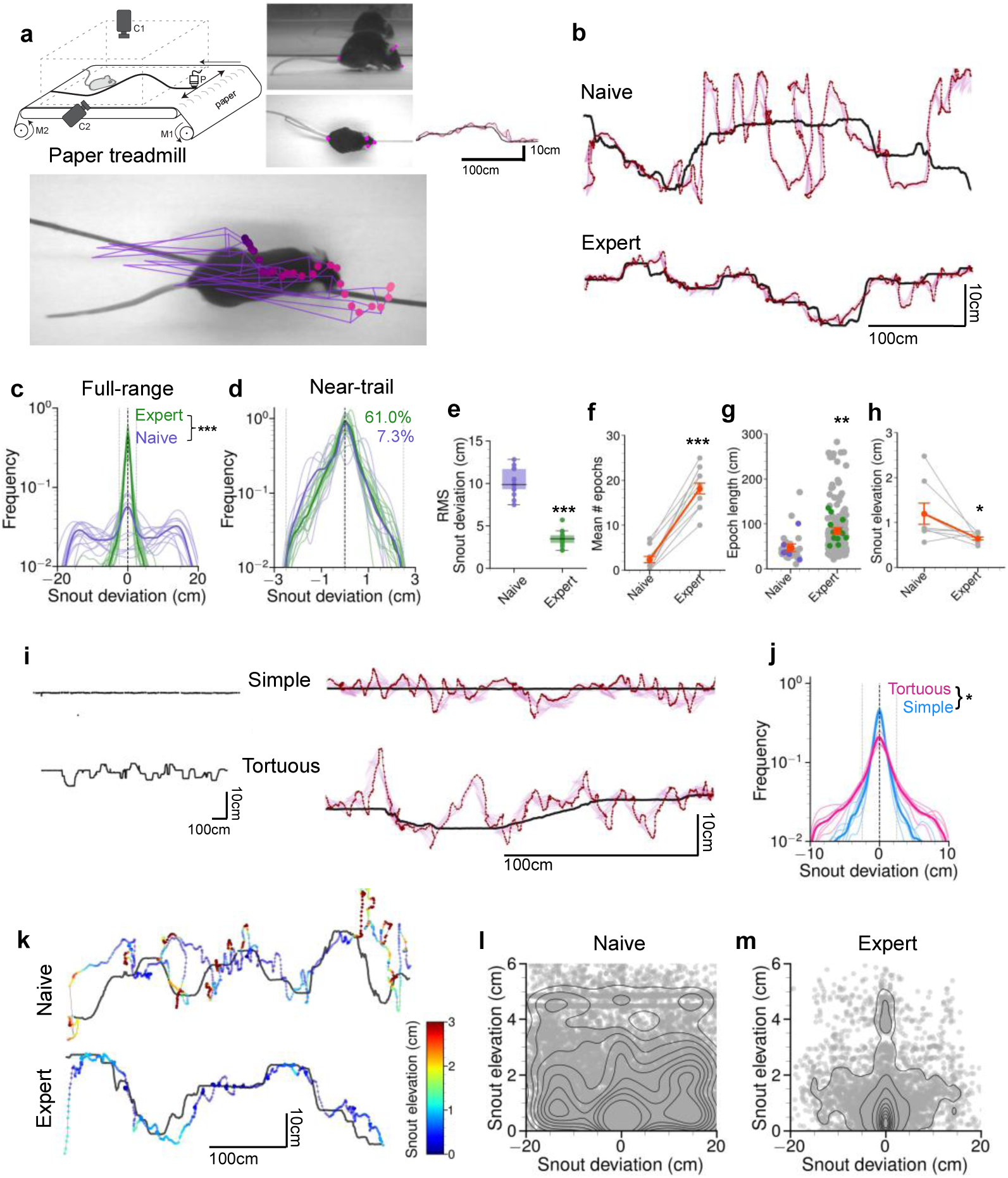
Mice rapidly learn to follow continuous trails. **(a)** Schematic of the custom-built paper-based treadmill. M1 & M2: rollers, with M1 coupled to a DC motor; C1 & C2: cameras; P: printhead that houses the ink with odor. The mice are restricted to the width of the chamber with walls made of clear acrylic that allow imaging with cameras. Odors are delivered using a printhead or a brush, controlled by stepper motors, allowing us to deliver varying geometries on the fly. Key points on the mouse (snout, right and left ear bases as well as base of the tail) identified using DeepLabCut (Mathis et al., 2018) are shown in magenta. **b)** Examples showing the trajectory of a mouse on day 1 (naïve) and on day 6 (expert) following a trail. The trail is shown in black. The snout is shown in purple, with a rhombus joining the snout, ears and base of the tail in pink. **c)** Distributions showing the snout deviation from 13 mice when naïve (day 1) vs when they have become experts (day 6), D =0.32 **,\*\*\***p<0.001, Kolmogorov–Smirnov test. Each line indicates a unique animal. **d)** The same data shown in panel c, but subsampled to include only points closer than 2.5cm on either side of the trail, numbers indicate percentage tracking. (**e-h)** Performance metrics of following a trail in naïve and expert mice. Each dot represents an individual animal, except in g where each colored dot represents an animal. **(e)** Root mean square deviations from the trail either in naïve or expert animals. **\*\*\***p<0.001, Two-tailed Wilcoxon rank-sum test. **(f)** Mean number of epochs, **\*\*\***p<0.001, Two-tailed Wilcoxon rank-sum test. **(g)** Length of each epoch, **\*\*\***p<0.001, Mann-Whitney U test and **(h)** Elevation of the snout from the paper floor, **\***p<0.05, Two-tailed Wilcoxon rank-sum test. **(i)** Left: 2 example trails presented to mice. Right: Examples of a mouse following either a simple straight or a tortuous trail. **(j)** Distributions of snout deviations for 5 mice when following either a simple or a tortuous trail. **(k)** Examples of trail tracking in naïve or expert mice, with the vertical elevation of the snout mapped on to the tracks (colormap representing snout elevation shown to the right). **(l, m)** Distributions showing snout occupancy in 3d space in naïve mice **(l)** and when they have become experts **(m).**

We could influence how mice sample the odor by varying the speed of the paper in the arena. We found that trail tracking fidelity varies as a function of the trail width and paper speed (Figure S2a,b). However, the identity of the odor used had no effect on the fidelity of tracking (Figure S2c). Mice quickly learned to consume the rewards on the trail and by day 5 were able to collect around 70% of the stochastically presented rewards (Figure S2d,e). When tested with similar treadmill speed and reward statistics but without a trail, mice manage to collect < 30% of the overall rewards (Figure S2e), indicating that mice must follow the odor trail faithfully to obtain rewards at a high rate. In further support of this idea, mice collected significantly fewer rewards if the droplets were deposited away from the trail (Figure S2e).

By flexibly varying the position of the odor printer over time, we could present trails of diverse shapes. Mice that were experts at following simple trails were much less accurate when presented with tortuous trails with frequent kinks (Figure 1i, j, S2f,g,h) (52.0 ± 5.62% tracking in expert mice tracking simple trails as opposed to 16.8 ± 2.23%, mean ± s.e.m, when presented tortuous trails, (5 animals) p = 0.030, KS test). Conversely, mice trained only on tortuous trails quickly adapt to straight trails and track them accurately (Figure S2i) (28.2 ± 7.68% tracking percent in expert mice tracking tortuous trails vs 68.4 ± 6.53% when switched to straight trails, mean ± s.e.m, 4 mice, p = 0.125, Wilcoxon signed-rank test). Mice lower their snouts to the ground to sample the trail, presumably to maximize the signal from volatile odors that disperse quickly above the ground. We used triangulation from multiple cameras to characterize their snout movements in three-dimensional space (see Methods). The vertical distance from the snout to the paper was highly variable on the first day and became tightly distributed around the trail over a few days, indicating that mice start sampling the ground more than the air as they become better at following odor trails (Figure 1k-m). When the trail odor was also presented as an airborne cue, expert mice are poorer at following the trails, indicating that mice track the odor and not some texture or scent from the printer ink (Figure S3a, b).

Overall, our experiments show that mice naturally follow odor trails but learn to improve their tracking fidelity over multiple days, and that trail tracking accuracy varies with the geometry of the trails encountered and the speed of the treadmill.

### Bilateral inputs are helpful for accurately tracking odor trails

What are the strategies used by mice to track odor trails? Prior studies have suggested that bilateral signals arriving through the two nostrils aid in odor-guided navigation, including tracking an odor trail (Catania, 2013; Jones and Urban, 2018; Khan et al., 2012; Rabell et al., 2017; Rajan et al., 2006). These studies offered evidence for a role of bilateral olfaction using selective blockade of one nostril. After mice became experts at tracking odor trails in our set up, we stitched one nostril (9 mice for left and 6 mice for right) and monitored how they followed trails. Consistent with previous studies, we observed that mice with just one intact nostril were still able to follow trails, but with a lateral bias (Figure 2a,b;D = 0.12, p = 0.0009, KS test, left nostril compared to sham). The biases were on the same side of the stitched nostril (Figure 2b, c and Figure S4a-d, lateral signed bias of 0.007 ± 0.02 cm in sham, compared to 0.074 ± 0.02 cm with the right nostril stitched, −0.096 ± 0.04 cm with the left nostril stitched, mean ± s.e.m.), such that mice placed their good nostril close to the trail and the closed nostril away from the trail. We also found that the number of tracking epochs and the length of each epoch decreased in nostril-stitched animals, whereas the snout elevation was not significantly different from control (Figure S4e-h, 18.7 ± 1.64 epochs in sham as opposed to 13 ± 1.32 epochs, mean ± s.e.m. when the nostril was stitched, p = 0.02135, Mann-Whitney U test). The biased tracking persists even after a week following nostril stitching (Figure S4i, 4 mice).

**Figure 2:**
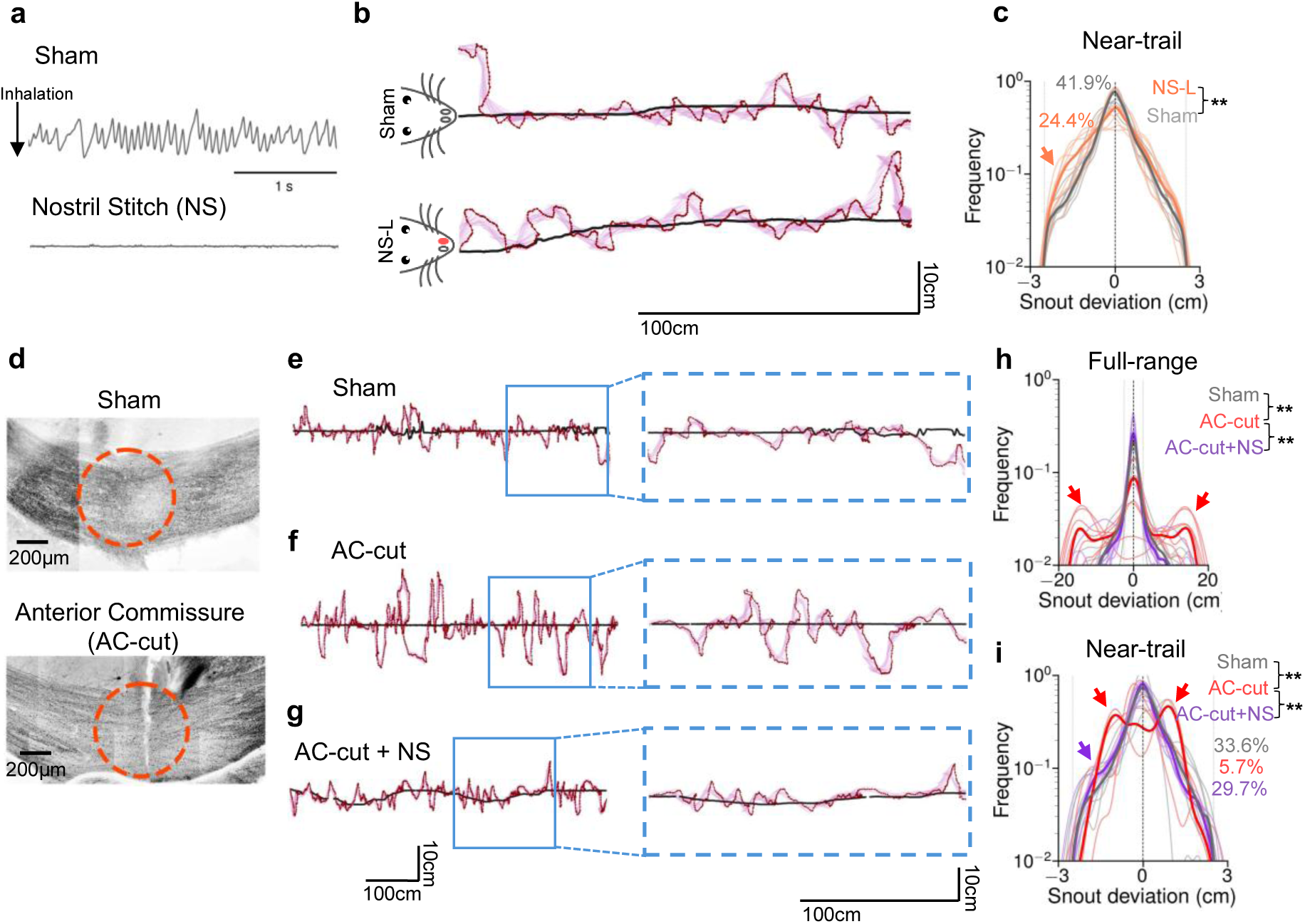
Coordination of bilateral inputs is helpful for accurately following trails. **(a)** Example time series from a thermocouple in a sham operated and a nostril stitched mouse. The thermocouple signal was at zero when the corresponding nostril was blocked. The arrow indicates inhalation. This was used to confirm that the nostril occlusion prevented airflow on that side. **(b)** Example trajectory from a mouse following a trail either after a sham stitch or when the left nostril was stitched. **(c)** Distributions of the deviations of the snout from the trail from either mice where there was a sham stitch or when the left nostril was stitched,D = 0.12 **\*\*\***p<0.001, Kolmogorov–Smirnov test, numbers indicate percentage tracking, p<0.01, Mann-Whitney U test. Orange arrow indicates the bias toward the closed side. **(d)** Histological evidence for the transection of the anterior commissure. **(e-g)** Example trajectory from a mouse with either sham surgery, one with the anterior commissure transected and with an animal where the nostril was stitched post severing the anterior commissure. Data is aligned to the perturbed nostril. Inset shows zoomed in sections. **(h,i)** Distributions of snout deviations from at least 5 mice in conditions shown in (e,f,g) for all periods of behavior **(h)** or selected periods when mice were within 3cm of the trail **(i)**,**\*\***p<0.01, Mann-Whitney U test. Numbers indicate percentage tracking. Red arrows indicate the bias due to commissure cut, while purple arrow indicates the bias due to the stitched nostril and a severed commissure.

Closing one nostril not only creates an asymmetry in stimulus acquisition but also likely reduces the total signal acquired. Additionally, this perturbation could lead to compensatory alteration of the strategies used for following trails, which might mask those used in unperturbed conditions. Therefore, we turned to an alternate manipulation that leaves bilateral stimulus acquisition intact but interferes with communication between the two streams. After mice had become expert at tracking trails, we severed the anterior commissure, the main fiber bundle that connects the ventral olfactory regions in the two hemispheres, in particular the AON and piriform cortex (Brodal, 1948; Brunjes, 2012; Gurdjian, 1925; Kikuta et al., 2008; Schoenfeld and Macrides, 1984) (Figure 2d). Mice with severed anterior commissure follow trails poorly, taking more tortuous paths (Figure 2e, f). The number of tracking epochs and their duration as well as the snout elevation were significantly different upon cutting the commissure (Figure S5a-d; 15.7± 1.48 epochs in sham (5 animals), 2.6 ± 1.07 epochs, mean ± s.e.m., when the commissure was severed (7 animals), p= 0.0011, Mann-Whitney U test). Interestingly, mice with their commissures transected failed to improve much over time (Figure S5e).

Careful inspection of the distributions of the snout deviations suggests that mice whose commissure was cut displayed a lateral bias toward one side of the trail or the other (Figure 2h). This suggested that there could be a form of rivalry between the two independent nostril inputs that remain unresolved in the absence of interhemispheric communication. To test whether there was such a rivalry, we stitched the nostril of the mice with their commissures cut. We observed that mice with their commissure cut, but with only one functional nostril, were able to follow an odor trail much more accurately, comparable to sham operated mice (Figure 2g-i, 22.7 ± 5.32 epochs when the anterior commissure was severed and one nostril blocked (5 animals) vs 15.7 ± 1.48 epochs in sham operated, mean ± s.e.m, p > 0.6, Mann-Whitney U test). This double manipulation also indicated that the commissure cut does not lead to some non-specific defects that cause animals to follow trails poorly. The behavioral rescue upon shutting one nostril is lost upon removal of the stitch.

These data indicate that, while mice can follow trails with a single intact nostril, bilateral information flow needs to be coordinated via the anterior commissure when both nostrils receive odor inputs.

### Mice turn reactively after asymmetric trail encounters

Mice predominantly acquire information about the odor trail when they sniff (Findley et al., 2021; Kepecs et al., 2006; Wesson et al., 2009). To determine whether and how mice modulate sniffing to aid efficient sampling and tracking, we measured respiration in mice following odor trails either using a thermocouple or using wireless diaphragmatic EMGs (Figure S6a-c). Wireless recordings had the important advantage of allowing free head movements, which may be difficult with tethers necessary for other methods.

Mice displayed high rates of respiration when following the trail, in comparison to much lower rates when they were further away from the trail (Figure 3a,b). The respiration rate dropped to low levels when mice were consuming the rewards presented sporadically on the trail, in line with work on coordination of various orofacial rhythms that report low rate of breathing when swallowing (Kaku et al., 2025; Kleinfeld et al., 2023; Moore et al., 2013; Saito et al., 2003).

**Figure 3:**
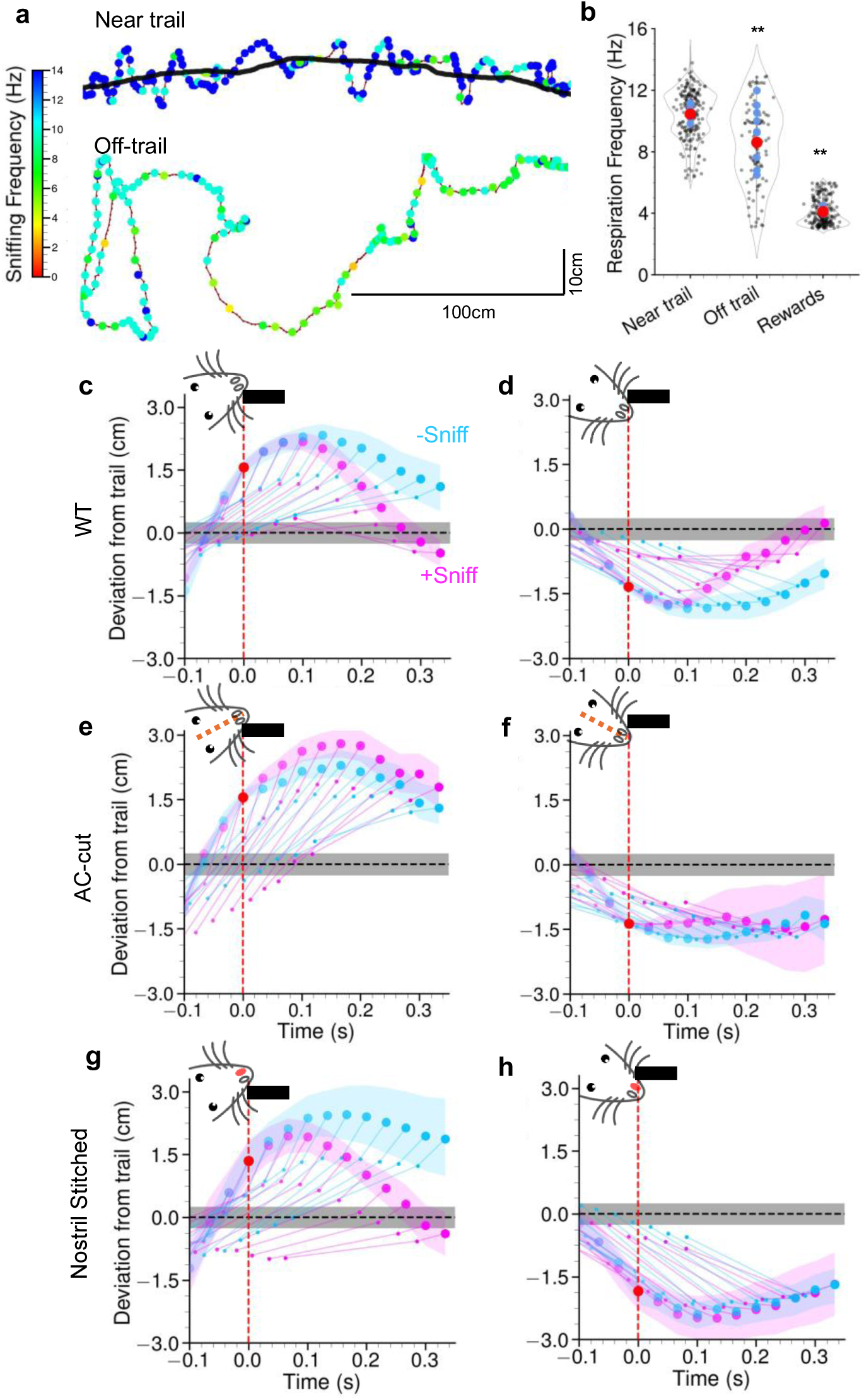
Modulation of respiration during trail tracking allows quick turns at trail edges. **(a)** Example trajectory of a mouse either following a trail or when there is no trail. Inhalations are shown as colored dots. The colormap shows instantaneous respiration rate. **(b)** Average respiration rates in different conditions, **\*\***p<0.01, Kruskal-Wallis ANOVA with post hoc Mann-Whitney U tests using Bonferroni multiple comparisons correction. **(c-h)** Average mouse trajectories showing snout (large circles), shoulder and base of the tail, joined by lines. The red dot indicates the anchor point for the averages, when the mouse snout was within 1 - 2cm of the trail after crossing it. Pink traces are averages when an inhalation was detected at t = 0, and cyan are when there is no inhalation at t = 0. Left and right panels are separate averages for trail crossings in opposite directions. Trail of 0.5cm is shown as a gray line. **(c, d)** is from wild type mice (at least 27 stretches from 12 mice) and **(e, f)** shows data from mice where the anterior commissure was transected (at least 28 stretches from 8 mice). **(g, h)** shows data from mice where the left nostril was stitched (at least 25 stretches from 5 mice).

Previous studies (Catania, 2013; Jones and Urban, 2018; Khan et al., 2012), as well as our experiments with nostril occlusion and commissurectomy, suggest that mice may use bilateral comparison of odor intensity to track trails efficiently. We searched for signatures of such a computation in our data by asking whether asymmetric intensity signals across the two nostrils can trigger an orientation response. We computed the average trajectory of the snout and head of the mouse conditioned on whether or not there was a sniff when the snout was located at specific distances from the trail, from at least 25 stretches from 12 animals. An inhalation that occurs when the mouse is around 1.5 cm from the center of the trail as it is moving away from the trail triggers a reorientation of the head towards the trail within around 150 ms (Figure 3c,d). The head reorientation was followed by a full body turn. This rapid reorientation occurs much less prominently when there is no inhalation. Importantly, this reorientation does not occur after inhalation when the snout is much closer to the trail, such that the two nostrils are more symmetrically placed on the trail (presumably leading to similar odor intensities reaching the two nostrils) (Figure S6d,e). In addition, the distance at which the reorientation occurs varies as a function of the trail width. An inhalation triggers rapid reorientation closer to the trail with a narrower trail (Figure S6f-i).

If the reorientation is triggered by computations using the asymmetric sensory inputs to the two nostrils, disrupting the comparison of the two inputs should prevent this behavior. To test this idea, we repeated this analysis in mice with severed anterior commissure and found that there was no privileged reorientation triggered by inhalation (Figure 3e,f). We also found that the reorienting behavior was preserved in nostril stitched animals when the open nostril was closer to the trail, but impaired when the closed nostril was closer to the trail (Figure 3g,h). Under conditions where the nostril was stitched, the differential reorientation occurred further away from the trail (Figure S6j,k).

Collectively, these analyses suggest that upon inhalation, if there are differences in odor intensities in the two nostrils, mice orient towards the trail with the intensity comparison requiring interhemispheric communication through the commissure.

### Mice use past trail encounters to estimate future heading

The reorientations triggered upon inhalation described above indicate that mice can use reactive strategies during trail tracking. It remains unknown whether and how mice use knowledge about trail features, internalized from prior experiences (long-term memory) or recent experiences (working memory or adaptation) to make decisions about their actions. Earlier experiments have indicated that rats cast with increasing amplitude upon losing the odor trail (Khan et al., 2012), which could be due to a simple expectation of the trail based on the preceding sniff. A recent theoretical model formulated a sector search strategy that is based on a short-term memory of accumulated information from previous samples (which could result in presence or absence of odor) (Reddy et al., 2022b). We performed three separate experiments to test whether mice use recent experience to follow trails.

First, we presented long straight trails, which were then interrupted by breaks. Upon encountering breaks mice perform casts of increasing amplitude (Figure 4a,b). Breaks introduced within complex tortuous trails lead to more rapidly growing casts by mice (Figure 4a,c), likely reflecting memory of recent trail characteristics and greater uncertainty in their trail heading estimates.

**Figure 4:**
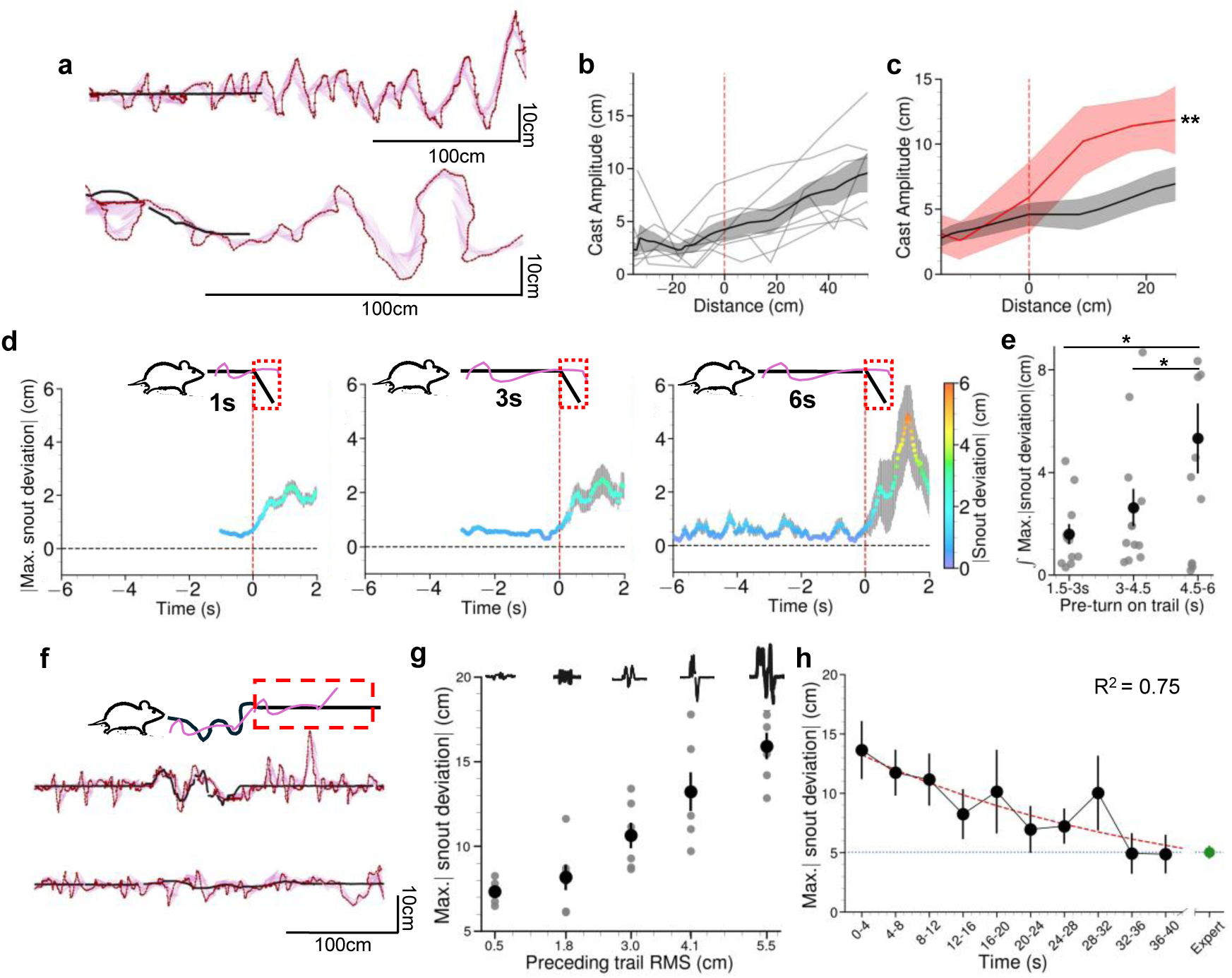
Mice use memory of previously encountered trail statistics. **(a)** Example trajectories from mice encountering a break while tracking a trail that is either straight (top) or tortuous (bottom). **(b)** Casting amplitude from 7 stretches from 5 mice encountering a break when following a straight trail. **(c)** Casting amplitude from 7 stretches from 5 mice encountering a break either when following a straight trail (black) or a more tortuous trail (red). **(d)** Snout deviations over time as mice encounter a sudden turn while following a trail. **(e)** Averages of the integrated maximum snout deviations over 2 seconds after encountering a turn after different periods of following a straight trail. (*p<0.05, Kruskal-Wallis ANOVA with post hoc Mann-Whitney U tests using Bonferroni multiple comparisons correction). Each point is for one session from one mouse. **(f)** Example trajectories from straight sections of a trail before or after encountering noise patterns in the trail **(g)** Maximum snout deviation in sections where the trail was straight preceded by trails of varying noise patterns. **(h)** Maximum snout deviation in a straight section of the trail immediately after encountering a trail with a root mean squared deviation of 4.1cm. The average maximum snout deviation from 9 expert mice that have not faced noisy trails are shown in green. The R^2^ value indicates goodness of fit, p-value=0.9032 (Chi-squared test). The best fitting time constant was 18.4 seconds.

In a second set of experiments, we introduced unexpected sharp turns in the trail after varying periods of straight stretches. Mice make excursions away from the trail upon encountering sharp turns (Figure 4d, Figure S7a-c). If mice adapt to straight stretches, they might predict the future trail geometry to also be straight. If so, the length of experience in straight stretches would influence their behavior when encountering the unexpected turn. Indeed, when mice reach the turn after longer periods of tracking straight trails, they overshoot by a larger amount before correcting their heading (Figure 4e). This overshoot was not due to differences in forward velocity of the mice when reaching the turn (Figure S7d,e).

A third set of experiments involved the presentation of straight trails interspersed with periods of convoluted trails with different degrees of lateral excursions. In this design, mice will arrive at the straight stretches after periods of following trails with different convolutions. Remarkably, mice follow highly similar straight trails with very different accuracies, exhibiting much greater lateral deviations in the straight sections after following trails that are more convoluted (Figure 4f, g). This dependence of lateral deviation on the tortuosity of preceding trails was also seen in nostril stitched animals (Figure S7f). To test how long the altered pattern of trail tracking persisted, mice were presented with one version of the convoluted stretch (with a root mean square deviation of 4.1cm) for 9-11 seconds, followed by a long straight segment. This experiment indicated that the altered trail tracking pattern returns to the baseline pattern of tracking a straight trail with a time constant of 18.4 seconds (Figure 4h).

Taken together, these findings suggest that mice use an internal model of recently encountered trail statistics to estimate the trail heading direction and uncertainty, which informs the actions taken to follow the current trail.

### Trail tracking models suggest extended memory is required for accurate predictions

Predictive search strategies use the history of past trail detections and internal models of the tracked trails to make inferences of their future heading. For instance, the animal may infer the future heading and its associated uncertainty to define an angular sector within which it is most likely to find the trail (Reddy et al., 2022b). Unveiling the details of the internal models used by mice is a major goal that extends well beyond the scope of our work here, but we can use the above data to compare simple reactive and predictive models. First, we reproduced the model from Khan et. al. (Khan et al., 2012), which imposed automatic lateral movements of the nose, with turns that were modulated by spatial (across two nostrils) or temporal (across two sniffs) odor gradients. An internal estimate of the trail heading and error bounds were calculated with hand-crafted parameters that exploit up to the past five detections. Therefore, the model has a limited memory with some heuristic “turning back” triggered by the nose going too far out of the heading estimate. An agent with these features and parameters can track fictive odor trails of simpler geometries similar to a purely reactive model (Figure S8a-d,i) without memory but fails to follow trails with gaps and is much worse on more complex trails with higher tortuosity (Figure S8e-h,j). The model also fails to recapitulate the history-dependent effects we observed in Figure 4d-g.

Therefore, for the agent to track efficiently under more difficult conditions, a reactive strategy alone is not sufficient, and a more extended temporal memory is needed. A Bayesian formulation of the problem of inferring the future trail heading (the posterior density) based on the current measurement (likelihood) and the previous accumulated knowledge of the trail from the past observations (prior) reads as follows

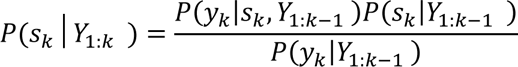

Here, *Y*_1:*k*_ is the set of all detections from the beginning of the hypothesized memory length up to the current time, *i.e.*, *Y*_1:*k*_ = (*Y*_1:*k*−1_, *y*_*k*_) with *y*_*k*_ denoting the last measurement (sniff-related trail odor detection or non-detection). The variable *s*_*k*_ denotes the inferred state of the future trail heading, which could involve various statistics, such as mean heading, its uncertainty, curvature of the trail, frequency of breaks, etc. The first term in the numerator is the likelihood of a detection/non-detection, while the second is the inferred state of the trail prior to the current measurement. The denominator is a normalizing factor corresponding to the overall evidence. The goal of an agent tracking the trail is to estimate this posterior density (the left-hand term in the above equation) at each instance of decision making, which can be discretized at some acceptable resolution, and control its motion accordingly. A purely reactive strategy will rely entirely on the current measurement, *i.e.*, *P*(*s*_*k*_│*Y*_1:*k*_) = *P*(*s*_*k*_│*y*_*k*_).

The use of prior knowledge is a key ingredient of classic state estimation procedures, such as Kalman filtering or the more general Bayesian particle filtering (Arulampalam et al., 2002; Chen, 2003; Särkkä and Svensson, 2023). Previous models of olfactory navigation have postulated an extended memory (Vergassola et al., 2007), or a more limited one (Khan et al., 2012; Reddy et al., 2022b), but it remains unclear whether and how mice (and other animals) use recent experience to obtain these estimates. In order to highlight the diversity of possible solutions to the problem, and since the form of the prior is unknown, we used Gaussian Process Regression (GPR) to estimate it with minimal assumptions. (Arulampalam et al., 2002; Rasmussen and Williams, 2005) (see Methods). Our phenomenological model is meant to illustrate how internal beliefs derived from noisy and limited memory lead to estimates of future trail heading that drive behavior over timescales relevant to real navigation.

An agent that used GPR to estimate the prior successfully tracked trails of varying geometry (Figure 5a-e). The agent was programmed to: (1) turn towards the most likely heading upon encounter and update the heading estimate uncertainty (grey shaded regions), (2) reverse direction when hitting the boundary of the uncertainty estimate. We estimated the most likely heading and the associated uncertainty using varying histories. Since it was unrealistic to assume animals weigh every contact equally, which would require infinite memory extending backward in time, we used an exponential temporal weighting scheme to bias the GP fit toward more recent crossings, using *w*_*i*_ = exp (−*λ*(*t*_*last*_ − *t*_*i*_)), where *w*_*i*_ is the weight assigned to the *i*-th crossing and the *t*_*last*_ − *t*_*i*_ is the time since last trail crossing. Varying *λ* changes the recency bias. Estimates from agent-based simulations suggested that taking *λ* = 0.5 resulted in a memory of about 3.4 ± 0.49 seconds, whereas taking *λ* = 0.05, resulted in a memory of about 15 seconds (Figure S9a). Interestingly, deviation measurements from the trail from simulations suggest that a memory of about 15.8 ± 0.91 seconds performs significantly better than that of about 3 seconds (deviation of 1.72 ± 0.13 cm as compared to 1.28 ± 0.12 cm; Figure S9b-f). Further increase in memory does not produce significant improvement in tracking a tortuous trail (Figure S9b). This preferred time scale is remarkably similar to the history estimate for mice (Figure 4h).

**Figure 5:**
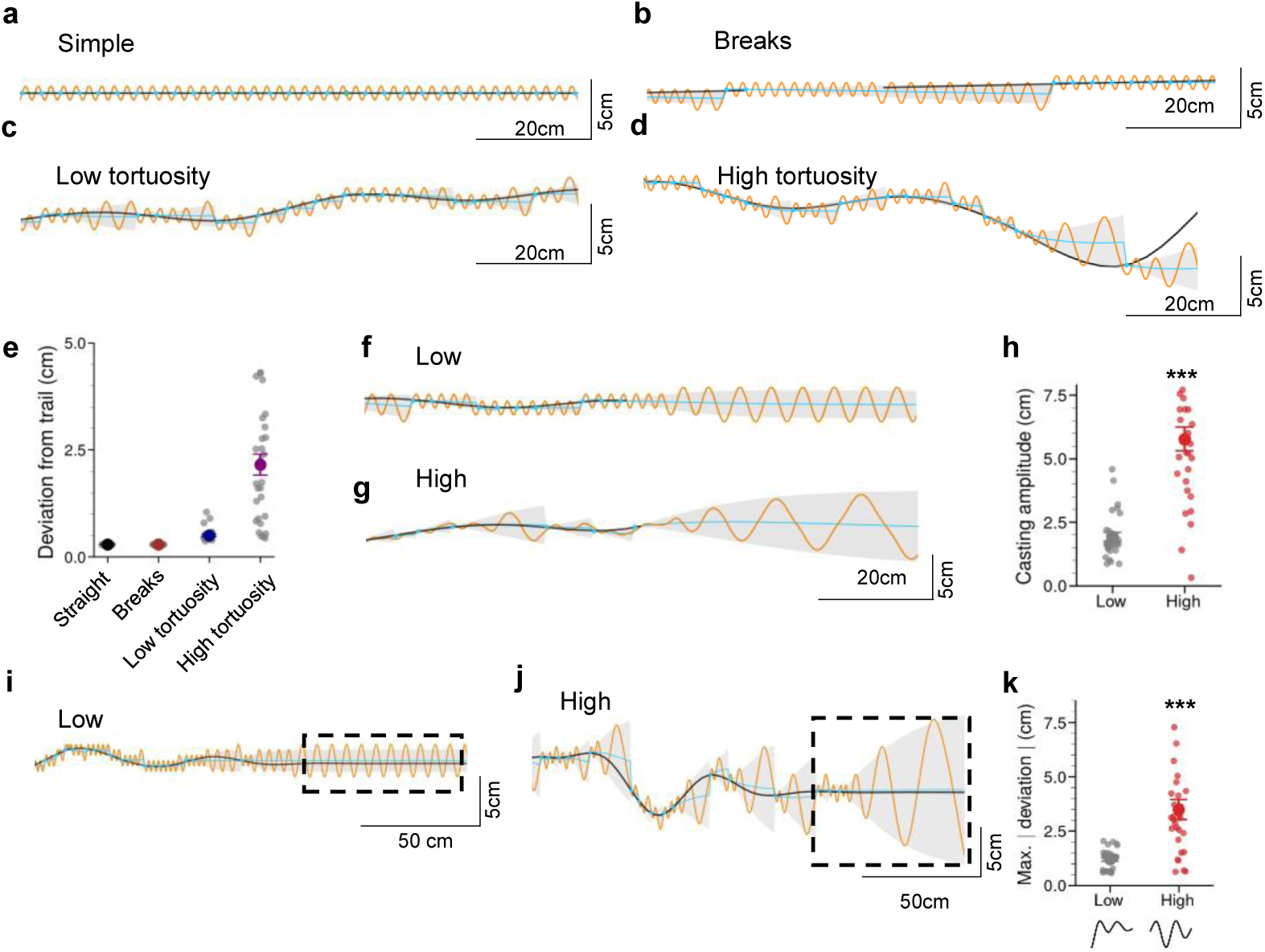
An agent using memory reproduces many facets of tracking behavior. **(a-d)** Example trajectories of an agent using Gaussian Process Regression to estimate the trail from previous contacts with the trail. The virtual trail is shown in black, the agent’s trajectory in orange, the agent’s trail encounters as sky blue dots, the agent’s heading estimate in a sky blue line and the posterior uncertainty of the trail position (± 2σ, where σ is the standard deviation output by the GPR at each point) in gray cones. The example trajectories are from the agent following **a)** a simple straight line, **b)** a simple trail with breaks, **c)** curves of low tortuosity and **d)** curves of high tortuosity. **(e)** Deviations from the trail as shown by means ± s.e.m. are from 30 simulations, show that the agent is accurate in following trails. **(f** and **g)** Example trajectories of the agent encountering a break while following trails of two types of curvature. **(h)** Casting amplitudes measured from 30 simulations as shown by means ± s.e.m. show that the agent casts more widely when encountering a break following the trail of higher curvature. **\*\*\***p<0.001, Mann-Whitney U test. **(i** and **j)** Example trajectories from an agent following a straight section of a trail after encountering either low tortuosity **(i)** or high tortuosity **(j)**. **(k)** Snout deviations over time from agents following a trail. The deviations are from straight sections of the trail. The deviations are significantly higher when the trail was preceded by the curves of higher tortuosity. \***\*\***p<0.001, Mann-Whitney U test.

We next tested the behavior of the agent upon encountering a break after having followed trails of either low or high tortuosity (Figure 5f-h). We find that the agent’s heading is more uncertain when the break is preceded by higher tortuosity (Figure 5g compared to Figure 5f) and is reflected in the increasing casting amplitude (Figure 5h), resembling what we observed in mice (Figure 4a-c). Finally, we examined the effect of an agent with extended memory on trails that straightened after being convoluted as in Figure 4f. The agent qualitatively replicated the behavior of mice (Figure 5i-k). The trail tracking in the straight stretch has more lateral deviation when the agent has just been following a convoluted trail, since the uncertainty estimate continues to be influenced by the past experience within the memory window. The more convoluted the prior trail, the greater the lateral deviation in the straight stretch (Figure 5j, k).

To explore alternative predictive models to explain our data, we adapted the sector search framework (Reddy et al., 2022b) which used a Generalized Worm-Like Chain (GWLC) prior to inform a Bayesian agent that tracks trails using local trail heading and uncertainty from recent trail contacts. In the GWLC setting, we replaced the two-contact heading estimate used in that earlier work with a principled inference from a finite history of trail encounters, thereby endowing the sector orientation and aperture with an explicit, tunable memory for recent tortuosity (Figure S10a-l). We also considered an Extended Kalman Filter (EKF)-based Bayesian agent, which compresses past encounters into a latent state and covariance (Figure S11a-l). Both agents reproduced the dependence of heading uncertainty and casting amplitude on prior tortuosity (Figure S10e-k, S11e-k), indicating that a finite memory for recent trail structure is sufficient to generate key aspects of the behavior observed in mice (Figure 4e and g). In addition, the GWLC formulation captured the increased lateral deviation on straight segments following convoluted stretches. Interestingly, the GWLC formulation also captured the recovery to baseline accuracy of tracking in the straight section (Figure S10k and l). The EKF agent captured the overshooting behavior in the straight section provided they reached the straight section (Figure S11k and l). These results point to an underlying principle where mice behave as if they carry forward a probabilistic summary of recent trail history, rather than reacting only to their most recent encounters.

Taken together, these results show that simple reactive or fixed rules fail to replicate key features of trail tracking observed in mice. In contrast, Bayesian filtering models which use recent trail history to dynamically estimate heading direction and associated uncertainty reproduce observed mouse behaviors remarkably well.

### Neural perturbation and recordings implicate AON in trail tracking

To characterize the sensory information entering the mouse brain, we imaged neural activity in the axons of olfactory sensory neurons (OSNs) converging onto the olfactory bulb (OB) glomeruli in mice tracking odor trails using head-mounted miniature microscopes, while measuring respiration simultaneously (Figure 6a). Clear changes in fluorescence intensity could be detected in multiple glomeruli, and transients were extracted using CaImAn (Giovannucci et al., 2019) (Figure 6b-d, see sample video). Visual inspection suggested that calcium transients were more frequent when the snout was close to the trail and the animal took a sniff (Figure 6b,b’; Figure S12a-c). Quantitative analysis of snout deviation conditioned on the detection of glomerular activity revealed that transients were highly likely when the snout was < 1 cm from the trail in expert animals, and the likelihood dropped off steeply to low levels past 3 cm (Figure 6e, Figure S12a,b). Importantly, the steepest gradient of this distribution is around 1.5cm from the center (Figure S12c), matching the distance at which mice were found to reorient their snout upon inhalation (Figure 3c,d). This distribution was shallower and of lower peak height in naïve animals (Figure 6e), suggesting a change in sampling strategies as animals learn to track well. Sniff-aligned averages revealed robust glomerular responses when the snout was around 1.5 cm from the trail (Figure S12d). We found that accurate binary classification of near versus far from the trail (threshold distance of 1.5 cm) could be achieved using the activity of just a few glomeruli (Figure 6f), confirming that the information about the position of the trail is present in the input to the brain for downstream circuits to act on.

**Figure 6:**
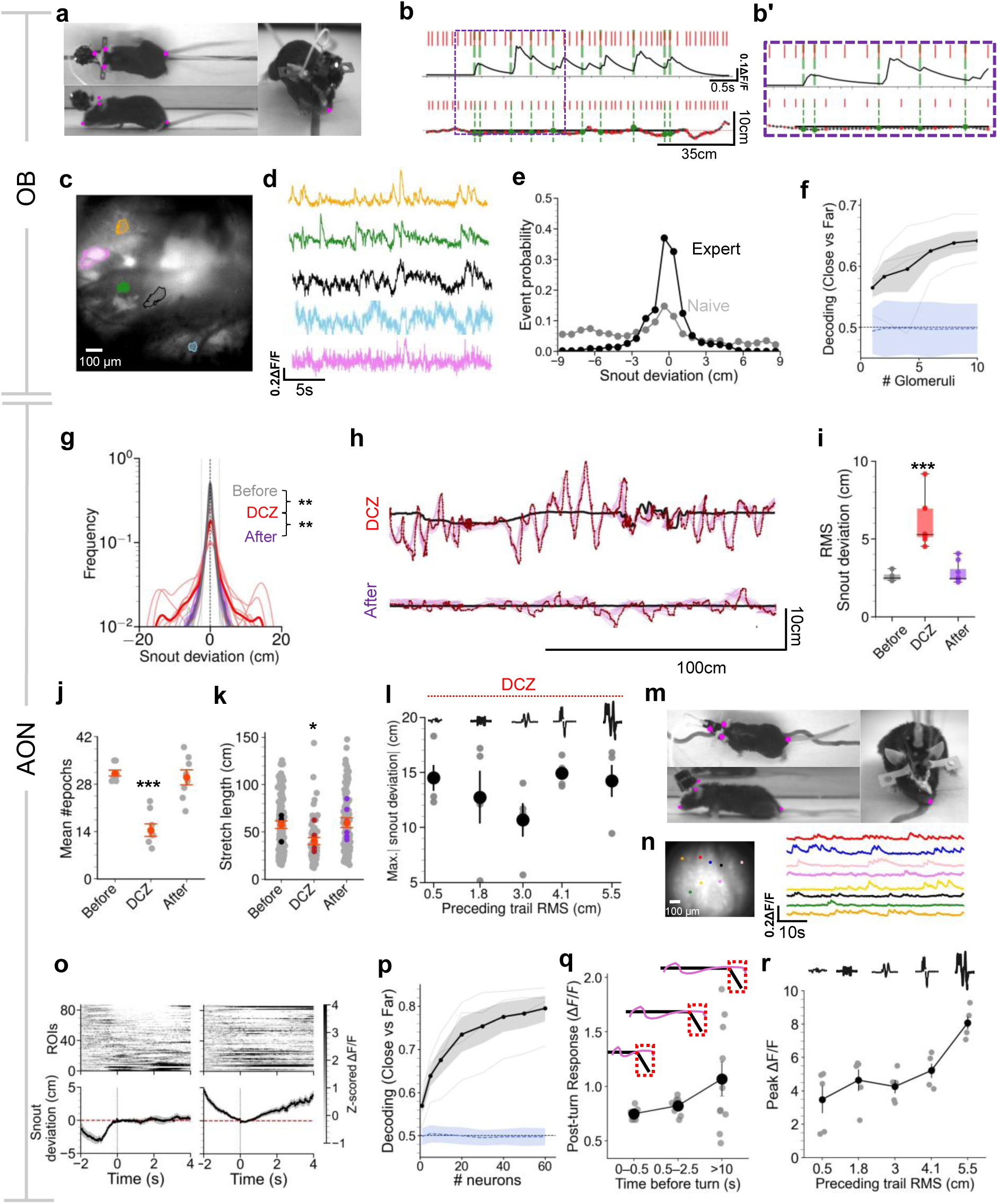
Neural representations during trail tracking behavior. **(a)** Views of the mouse with the head-mounted microscope recording from the glomeruli in a freely behaving mouse tracking a trail where respiration is also being recorded. **(b)** Example ΔF/F traces from an example glomerulus aligned to inhalation (green dashed lines). Box indicates a segment with an enlarged view shown in (**b’**). **(c)** Example field of view of olfactory bulb showing glomeruli with 5 example ROIs overlaid. **(d)** Example ΔF/F traces from 5 example glomeruli shown in (**c**). **(e)** Distribution snout deviation conditioned on glomerular response as a function of snout deviation for expert (black) and naive (gray) mice. **(f)** Decoding performance as a function of number of glomeruli (from 5 mice). Shuffled control shown in blue. (***p < 0.001, permutation test). **(g)** Distributions of snout deviations from 8 mice on the session before (grey), during (DCZ, red) and after (post-DCZ, purple) bilateral silencing of neurons in the AON using DREADDs. **\*\*\***p<0.001, Kolmogorov–Smirnov test. **(h)** Example trajectories with bilateral silencing using DCZ and the session with saline (post DCZ). **(i)** Root mean square deviation from the trail from either the day before, during or the day after neuronal silencing. (***p < 0.001, Kruskal–Wallis with post hoc test). **(j)** Mean number of epochs. (**\*\*\***p<0.001, Wilcoxon rank-sum test. **(k)** Length of each epoch, **\***p<0.05, Mann-Whitney U test. **(l)** Maximum snout deviation as a function of preceding trail RMS from 8 animals where the bilateral activity in the AON was perturbed using DREADD based inactivation. **(m)** Views of the mouse with the head-mounted microscope recording from the AON in a freely behaving mouse tracking a trail. **(n)** Example field of view and ΔF/F traces from multiple ROIs using a GRIN lens targeting the AON. **(o)** Population activity from the neurons in the AON from 1 session (top) and corresponding deviation of the snout from the trail (bottom) aligned to time points when the animal approached the trail from either side. **(p)** Decoding probability of distance to the trail as a function of number of neurons from the AON. **(q)** Post-turn response (peak ΔF/F in a 2s window after the turn) as a function of time where the trail was followed before encountering an unexpected turn from neurons in the AON. **(r)** Peak ΔF/F response as a function of current trail RMS from neurons in the AON. (Data is from 5 animals).

The anterior olfactory nucleus (AON) is one of the first cortical processing stations of odor information and has been implicated in bilateral integration (Grimaud et al., 2024; Kikuta et al., 2008; Rabell et al., 2017). To examine its role in odor trail tracking, we used chemogenetic inhibition using designer receptors exclusively activated by designer drugs (DREADDs) expressed bilaterally in excitatory neurons (Roth, 2016). Systemic administration of the ligand deschloroclozapine (DCZ) (Nagai et al., 2020), impaired trail tracking (Figure 6g-k). Snout deviation was significantly larger (2.60 ± 0.11 cm on the day before, 6.04 ± 0.55 cm on the day of DCZ and 2.81 ± 0.24 cm the day after), and the tracking bouts were less frequent and shorter (31.250 ± 0.861, the day before, 14.37 ± 1.86 the day of DCZ, and 30 ± 2.24 epochs the day after, mean ± s.e.m, 8 animals, p = 0.00067; with corresponding lengths per epoch 57.70 ± 4.04, 40.27 ± 3.78 the day of DCZ, 59.66 ± 5.1 cm mean ± s.e.m, 8 animals, p = 0.015, Kruskal-Wallis ANOVA with post hoc Mann-Whitney U tests using Bonferroni multiple comparisons correction) (Figure 6i-k). These impairments were specific (Figure S12e) and reversible, since mice return to expert performance in the following session (Figure 6i-k, Figure S12f). Unsurprisingly, mice following trails with AON activity perturbation did not exhibit history dependent tracking behavior (Figure 6l).

We next imaged neural activity in the AON neurons expressing GCaMP8m using head-mounted miniature microscopes while mice tracked trails, with concurrent recordings of respiration (Figure 6m,n, Figure S12g). Fluorescence changes in the AON robustly reflected sensory information (Figure 6o, Figure S12h), and trail proximity could be decoded from around 40 randomly chosen neurons, with accuracy increasing with the number of neurons (Figure 6p). The side of the snout relative to the trail was likewise decodable from AON population activity from around 20 neurons, even after controlling for distance from the trail (Figure S12i). We investigated whether non-sensory information, including signatures of predictive processing, was present in the activity of AON neurons. Remarkably, when mice encountered an unexpected turn after longer periods of straight tracking, activity in the AON population was greater, consistent with a larger prediction-error signal arising from stronger expectations of continued trail straightness. (Figure 6q, Figure S12j, (mean Spearman’s ρ = 0.68 across animals; Friedman test, p = 0.015). This increased activity was not due to increased respiration (Figure S12k). Moreover, the peak population-averaged AON activity of AON in the straight trail stretches was larger after periods of tracking more tortuous trails, again correlating with greater violation of predicted trail statistics (Figure 6r, Figure S12l). Collectively, these experiments identify AON as a key region involved in odor trail tracking, whose activity is consistent with representations of predicted trail structure and violations of those predictions.

## Discussion

In this study, we used a continuous paper treadmill to present mice with dynamic odor trails, allowing us to observe their behavior over extended distances and durations as they collected randomly placed rewards. Mice quickly learn to follow odor trails and become more accurate trackers over several days. While mice can follow trails with just one nostril, bilateral integration is critical for accurately following trails when both nostrils are intact. Mice quicken their respiration when following trails to increase sampling rates, and asymmetric placement of their nostrils when a sniff occurs triggers a head turn towards the trail, a response absent in commissure-transected animals. Importantly, tracking an odor trail involved more than immediate reactions: mice retained short-term memories of trail geometry, as seen in their larger overshoots at sharp turns following longer periods of straight trails, and their adaptation to different trail patterns. A Bayesian model for inferring trail properties using prior trail encounters with a memory lasting around 15 seconds could explain the mouse behavior well, whereas a purely reactive model or one with a simple fixed memory failed to explain the data. Neural activity in the sensory periphery revealed the sensing range of the mouse from the odor trail, which aligns well with the distances at which reactive reorientations towards the trail occur. Perturbation of activity in, as well as neural recordings from the AON implicate this region in predictive trail tracking behavior. These findings suggest that mice integrate real-time sensory input with internal representations to efficiently track odor trails.

Our study was inspired by previous work in rats (Khan et al., 2012), and offers significant new insights into trail tracking behavior. First, by using non-repeating trails, we avoided any memorization of the trail geometry by the animals. Instead, mice had to rely on the ongoing experience of the trail to guide their behavior. A second new finding was that mice can spontaneously track trails on the first day, and their improvement over days is mainly due to longer periods of engagement in tracking a trail as well as their 3D snout placement near the ground. Third, we show that nostril occlusion experiments, a mainstay of testing bilateral olfaction (Catania, 2013; Jones and Urban, 2018; Khan et al., 2012; Rajan et al., 2006), may have confounding effects that can be resolved by more nuanced manipulation of bilateral integration. Finally, by printing trails of many different geometries, we revealed previously unknown ability of mice to make inferences about the statistics of the trail. The ability to print long trails whose thickness, shape and integrity can be controlled dynamically, combined with modern markerless tracking and automated behavioral analysis (Mathis et al., 2018), facilitated the collection of extensive data under many different conditions. Such high throughput acquisition in a controlled laboratory environment allows a balance between stimulus control and presentation of highly variable, non-repeating segments that are hallmarks of naturalistic behavior.

One strategy proposed for detecting and remaining on the trail is bilateral comparison of odor intensities across the nostrils, which might help detect trail edges. Evidence for this idea has largely come from the effects of blocking one nostril. We confirm previous findings (Catania, 2013; Jones and Urban, 2018; Khan et al., 2012) that mice can still follow trails, but less accurately and with a lateral bias. This perturbation reduces significantly the overall information acquired by the animal and it is likely that animals adapt their navigational strategies to their new state. Therefore, we created a situation where mice acquire information from both nostrils, but the interhemispheric communication between the two channels of information was disrupted by severing the anterior commissure. Mice were very poor at tracking in this condition, likely because information from the two channels could potentially be different (and conflicting), and without comparison may result in erroneous decisions which would take the animal away from the trail. Furthermore, the behavior does not improve over time. We note that the behavioral deficit is not due to some irreversible loss of navigational or other non-specific issues, since blocking one nostril in commissure-cut animals partially reverses the behavioral deficit.

The bilateral comparison hypothesis was further supported by our analysis that revealed sniff-triggered head reorientation. Sniffs to one side of the trail within a short distance of around 1 cm trigger head reorientation towards the trail within 150 ms. The same sniff taken when the nostrils are much closer to the trail do not trigger reactive head turns, presumably because the odor intensity is very similar in both nostrils. Supporting such bilateral comparisons, we found that sniff-triggered reorientations were absent in commissure cut animals. Taken together, our data suggests that an inhalation that leads to asymmetric activation of the two nostrils is detected in the brain through bilateral comparison and leads to head reorientation. There is clear evidence for reorientation of the nares towards an odor source presented asymmetrically to head-restrained animals, which is thought to be mediated by collicular circuitry (Rabell et al., 2017). We predict similar neural circuits are involved in head reorientation in our behavior.

We found strong evidence that mice form a predictive model of the trail structure and do not track odor trails using a purely reactive strategy. First, how mice search for the trail upon encountering a break is influenced by the recent trail structure. If breaks occur after a period of linear (straight line) trail, mice search with a slowly expanding casting cone, as observed before in rats (Khan et al., 2012), and rationalized by the sector search theory (Reddy et al., 2022b). In contrast, if a break is encountered after a period of highly tortuous trail with unpredictable bends, mice search in a rapidly expanding casting cone. This dependence of the search sector angle on the previous trail statistics cannot be reconciled with a purely reactive strategy, which depends on current sampling. Instead, it is consistent with predictive processing, resulting from a greater uncertainty in the estimate of the trail heading direction (based on previous detections) when following a tortuous trail (Jayakumar and Murthy, 2022; Reddy et al., 2022b). A second line of evidence comes from introducing unexpected turns in the trail while mice were following a long linear trail. If mice were on the linear section for a brief period, they deviated only a little when reaching the bend in the trail before correcting themselves. However, the longer mice were on the linear section, the greater the deviation when reaching the bend. This suggests that mice form a moment-by-moment expectation of the trail heading, integrating their past experience. The longer they are on a linear trail, the more confident they are about the future trail heading, which presumably helps them track linear sections better, but causes greater error when the trail deviates significantly from linearity. This finding offers further evidence of the use of predictive strategies by mice to navigate. Finally, we asked whether higher order features of the trail could influence tracking strategies in mice. We watched mice follow straight trails after a period of following convoluted trails of different complexities. Remarkably, mice followed identical linear stretches of odor trails with varying degrees of lateral deviations, depending on their recent experience. When following highly tortuous trails, mice seem to expect trails to deviate from linearity and predictively make larger lateral movements, leading to less accurate tracking of linear trails. Such experience-dependent differences cannot be accounted for by simple reactive trail tracking strategies.

Predictive strategies for odor tracking have been linked to a short-term working memory of heading direction in *Drosophila*. When navigating along a plume with a well-defined boundary, flies exploit a directional memory built from their recent exits from the plume to navigate back toward it (Siliciano et al., 2025). Similarly, transient odor encounters can establish a persistent heading that guides subsequent movement (Kathman et al., 2024). Together, these studies suggest that flies hold a compact navigational variable in working memory, namely a remembered heading direction. These and related studies offer clear evidence of working memory, but suggest mechanisms that appear simpler than the search strategies we see in mice, which make use of uncertainty in heading direction estimate.

We used a Bayesian filtering framework to build agent-based models for predictive trail tracking. In the absence of detailed knowledge about the neural circuits involved in this behavior, we chose to frame the agent’s memory of the trail in a general way using Gaussian Process Regression (GPR). In such a formulation, the agent’s estimate of the trail is obtained from a finite number of previous trail encounters. At any moment, the agent has an estimate of the future trail heading and the uncertainty associated with it. This is closely related to the sector search strategy (Reddy et al., 2022b), which uses Bayesian filtering to maintain a posterior probability distribution over possible trail headings by integrating past detections and non-detection, and where the agent then searches within a sector with high posterior probability. Here, we only use a finite memory of trail encounters, to enable flexible interpolation and explicit uncertainty quantification. Although a model with reactive turns and simple handcrafted extrapolations allows tight tracking for simple trail geometries, it fails for more complex trails that mice can track well. An agent that remembers up to 20 previous detections (memory time constant of 10-20 seconds) can successfully track tortuous trails as well as those with breaks. Interestingly, such memory also leads to less tight tracking of straight trails immediately following adaptation to tortuous trails. We further introduced a Generalized Worm-Like Chain (GWLC) based agent that uses recent trail contacts to infer a probabilistic estimate of local trail geometry. Weighing recent contacts more strongly selectively preserves behaviorally relevant information within a limited temporal window, rather than storing a redundant, unbounded trace of the past. Complementarily, the Extended Kalman Filter (EKF) based agent compresses this history into a low-dimensional latent state and covariance, illustrating how an efficient, redundancy-reducing neural code could implement approximate Bayesian filtering over recent trail structure and thereby realize a finite but functionally meaningful “memory” for tortuosity. The ability of agent-based models with memory to describe the mouse performance much better than previous simpler models offers strong evidence for the existence of predictive strategies.

Why are these short-term adaptations to the trail geometry helpful to mice? In sensory systems, adaptation to the statistics of stimuli increases information transmission in a predictive manner (Attneave, 1954; Barlow, 1961; Laughlin, 1981; Manookin and Rieke, 2023). In our experiments, we observed adaptation in the complete sensorimotor behavior. After a significant period of experience with straight trails, mice might benefit from reinforcing the prediction of future linear heading to lessen computational load and increase tracking speed. Similarly, adapting to sinuous trails with more elaborate lateral movements might improve trail tracking efficiency without having to update trail heading as often. These adaptations, while helpful when the trail geometry remains unchanged, lead to more errors when the geometry abruptly changes, as in our experimental design. Such adaptation can be rationalized as building of internal models of the trail geometry to allow predictive strategies for more accurate tracking (Fiser et al., 2010; Kawato, 1999; Rao et al., 2023; Todorov and Jordan, 2002).

We offer the first glimpses into the neural underpinnings of the different features of trail tracking behavior we have uncovered. The sniff-triggered sensory representation of odor information from the two nostrils could be integrated in regions such as the AON and the piriform cortex (Kikuta et al., 2010, 2008; Rabell et al., 2017; Wilson, 1997). These signals could then drive nostril and head reorientation towards the trail through collicular pathways (Rabell et al., 2017), followed by locomotor adjustments. In support of this hypothesis, we find that input responses measured in the OB indicate a sensing range of around 1.5 cm, which matches the snout deviation from the trail that best triggers a reorientation. Population neural activity in the AON carries information about distance from the trail, as well as the position of the snout relative to the trail, presumably due to the rapid drop in the intensity of the odor stimulus away from the trail. Importantly, chemogenetic suppression of activity in the AON reversibly impairs trail tracking.

An important aspect of trail tracking behavior is the modulation of such turning responses by previous history of detections. How sequences of detections and blanks are combined in the brain to obtain estimates of trail heading (goal or heading direction), so that appropriate corrections to head orientation can be made, remains unclear. The brain regions that estimate the direction of travel must also adapt to the statistics of the trail. We find that AON responses to loss of trail (due to an unexpected turn) scales with the length of previous tracking, suggesting coding of predictions and their violations. Similarly, the activity of AON in straight stretches of trail is a function of the preceding history of trail statistics. While this might reflect the distinct motor patterns (Figure 4f-h), it is clearly non-sensory in nature since larger AON responses occur when snout deviations are larger (when prior trails are more tortuous). Therefore, AON activity represents motor, predictive or error variables. Our findings set the stage for further investigation of the predictive computations that implement efficient trail tracking.

## Materials and Methods

### Animals

78 adult Black6/J (JAX #000664) mice of both sexes were used in this study. All animals were obtained from Jackson Laboratories and maintained within Harvard University’s Biological Research Infrastructure (BRI). Mice were housed in an inverted 12 h light cycle and fed ad libitum. Within the BRI, mice were group housed at 22 ± 1 °C at 30-70% humidity. Animals were 2-8 months old at the time of the experiments. Each behavioral session represents a unique animal throughout the experiments described here and multiple sessions have not been combined for any condition. A single behavioral session lasted around 4-6 minutes, during which animals freely encountered continuously printed odor trails on the moving paper substrate. Tracking behavior improved gradually across days. Trail-tracking episodes were defined as periods during which animals remained within 2.5 cm of the trail for at least 2.5 s. The effect of varying these thresholds is shown in Figure S1. All experiments were performed in accordance with the guidelines set by the National Institutes of Health and approved by the Institutional Animal Care and Use Committee at Harvard University.

### Behavioral apparatus

Mice run freely on an ‘endless’ paper spool within a clear acrylic box (45cm x30cm) that extended close to the width of the paper roll (McMaster-Carr, 20565T13). Ink trails were delivered using an adapted inkjet printhead or delivered through a paint brush coupled to a stepper motor that was programmed to create movements using custom written code implemented on a PRJC Teensy 3.6 micro-controller (32 bit 180 MHz ARM Cortex-M4 processor). Video recordings were obtained using 940nm IR LEDs (LED Lightsworld), which is essentially invisible to mice. Images were recorded at 30 Hz by three FLIR Grasshopper 3, 4.1MP Mono USB3 Vision, CMOSIS CMV4000 cameras, one from above, one from the side and one from the left corner of the acrylic box. For all analysis in this paper, only videos from the top and side were used. The trail was imaged from outside the box using a FLIR BlackFly S, 3.1 MP, USB3, Sony IMX265 camera. The paper spool was driven by a powerful DC motor (ClearPath) and the displacement of the paper was obtained using a linear encoder coupled to a wheel placed on the paper. The control of the DC motor driving the treadmill was implemented on a separate control system run on a PJRC Teensy 3.2 micro-controller (32 bit 72MHz ARM Cortex-M4 processor). Ensure drops were delivered on the trail as rewards using a solenoid controlled by the Teensy controlling the ink delivery. The rewards were delivered across the entire width of the paper at random times, drawn from an exponential distribution. Data was acquired using Bonsai (v2.8.5, https://bonsai-rx.org/ (Lopes et al., 2015)).

### Respiration measurements

Respiration measurements were obtained through either wireless EMGs targeting the diaphragm which is the main inspiratory muscle, or wired thermistors that were targeted to the olfactory epithelium. For wireless EMG recordings, the targeting was adapted from a procedure previously described (Hérent et al., 2020; Reisert et al., 2014). Briefly, mice were anesthetized using an intraperitoneal injection of ketamine/xylazine mixture (100 mg/kg and 10 mg/kg, respectively). Mice were placed in supine position, the peritoneum was opened horizontally under the sternum, the sternum clamped to expose the diaphragm and a sterile parafilm was used to protect the upper part of the liver. The electrodes were then sutured in place on the diaphragm and the Stellar BTA-XS telemetry implants (Stellar Inc., Woodland Hills, CA) implant was placed in the subcutaneous pouch made close to the ventral abdominal area. The implant was sutured in place through the suture tab on the device to avoid movement. The implants were modified to have custom fine-wire electrodes to facilitate attachment to the diaphragm. Buprenorphine extended-release (3.25 mg/kg) (Ethiqa XR) and Carprofen (5 mg/kg) was administered subcutaneously for analgesia at the end of the surgery. In addition, Sulfatrim (80mg/kg) mixed with Ensure was provided to mice to prevent infections. The implants were calibrated to store the recordings in millivolts, and the sampling rate of 500 Hz was used. Data was recorded onto the device and streamed after every session. Thermistors were implanted following procedures described previously (Findley et al., 2021; McAfee et al., 2016). Briefly, mice were anesthetized, and lidocaine was applied before exposing the skull. The epithelium underlying the nasal bone was exposed by drilling a small hole and the bead of the thermistor was inserted halfway through. The thermistor was held in place, was sealed using Kwik-cast and the mill-max pins were secured on the skull using a custom, 3D printed half head bar. Ultralight flexible tethers were made by soldering two wires (Cooner wire, CZ 1187 wire, FEP Insulation 38AWG with 0.012″ diameter) and the signal was amplified using a custom-built amplifier. The signal was sampled at 3KHz.

### Nostril stitch

We closed one nostril of mice using 7–0 gauge MicroSutures (S-N718SP13). Mice were given lidocaine on the nostril and anesthetized using an intraperitoneal injection of ketamine/xylazine mixture (100 mg/kg and 10 mg/kg, respectively). The suture was pulled through the upper lip of the nostril, and the nostril was closed completely. The closure of the nostril was confirmed using a drop of water placed on the nostril, which was not aspirated. For sham stitches, the mice were anesthetized, and a suture was just looped and the nostril was not closed. Mice were administered extended-release buprenorphine (Ethiqa-XR) (3.25 mg/kg).

### Commissure cuts

The anterior commissure was severed adapting an aseptic procedure described previously (Rabell et al., 2017). Briefly, mice were anesthetized using ketamine/xylazine and placed on a stereotaxic apparatus. Following a craniotomy, mice were injected with 200nl AAV-CAG-GFP (Addgene # 37825-AAV8) into the AON (AP:-2.9, ML:-1.2, DV:- 2.7 from the brain surface), using a pulled glass pipette attached to a nanoinjector (MO-10, Narishige). The commissure was severed in a second surgical procedure. Following a craniotomy, sapphire blade lancets (World Precision Instruments) were used (AP: −0.25 to 0.25, ML:0, DV:-5) to sever the entire anterior commissure. Sham animals went through the whole surgical procedure except that the blade was only lowered to a DV of 4mm thus preventing transection. Extended-release buprenorphine (3.25 mg/kg) (Ethiqa XR) and Carprofen (5 mg/kg) was administered subcutaneously for analgesia at the end of the surgery.

### DREADD based inhibition

For expression of inhibitory DREADDs, *vGlut1-Cre* (JAX #037512) mice were anesthetized using ketamine/xylazine and placed on a stereotaxic apparatus. Following a craniotomy, mice were injected with 400nl AAV-hSyn-DIO-hM4D(Gi)-mCherry (Addgene # 44362-AAV8) or AAV-hSyn-DIO-mCherry (Addgene # 50459-AAV8) into the AON (AP:-2.9, ML:-1.2, DV:- 2.7 from the brain surface) bilaterally, using a pulled glass pipette attached to a nanoinjector (MO-10, Narishige). The skin was sutured using absorbable sutures and extended-release buprenorphine (3.25 mg/kg) (Ethiqa XR) and Carprofen (5 mg/kg) was administered subcutaneously for analgesia at the end of the surgery. After a gap of four weeks to ensure optimal expression, mice were habituated in the behavioral rig and were presented with odor trails. After confirming these mice track the trails well, mice were briefly anesthetized with isoflurane and 0.9% saline was administered for 3 days consecutively before injection of saline or Deschloroclozapine (DCZ) (HelloBio, catalog # HB9126). DCZ was dissolved in 0.9% saline and administered i.p. at a dose of 0.1 mg/kg for all behavioral experiments. Mice were run 30 minutes after injection of either DCZ or saline.

### Histology

To visualize the transection of the anterior commissure and confirm GRIN lens and viral targeting, mice were deeply anesthetized with a ketamine/xylazine mixture and perfused transcardially with 10 mL of PBS (pH 7.4) first, followed by 10-20 mL of 4% paraformaldehyde in 0.1 M phosphate buffered saline (pH 7.4). After the brains were removed, 70-um thick horizontal sections were made using a vibratome (Leica). Slices were then washed and mounted for confocal imaging with DAPI mounting media and imaged with a confocal microscope (LSM 980, Zeiss).

### Trail generation and trail tracking agents

Different models of an agent tracking an odor trail were implemented. Fictive odor trails were generated as piecewise continuous functions, constructed from a sequence of linear and random (normally distributed) segments. Trail curves were interpolated via cubic splines over a fixed number of points (40 for low tortuosity and 80 for higher tortuosity), and breaks in specific conditions were introduced. Five types of agents were simulated viz. 1) a completely reactive agent, 2) an agent with limited memory and using odor gradient information to make estimates of the trail, conceptually similar to the model presented in (Khan et al., 2012), 3) an agent using the most recent contacts with the trail to fit a Gaussian Process Regression model (GPR) 4) a Bayesian agent using a Generalized Worm-Like Chain (GWLC) model to estimate trail heading, and 5) an agent using an Extended Kalman Filter (EKF) to recursively estimate local trail position, slope and curvature. All agents operated within the same simulated odor-trail environment and shared identical locomotor and sensory dynamics unless otherwise stated. The trail itself was never analytically available to the agent and could only be sampled intermittently through local odor encounters during locomotion. Odor concentration was modeled as a Gaussian function of the distance from the trail, 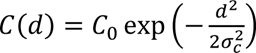, and odor detections were generated whenever the sampled concentration exceeded a fixed threshold. To avoid repeated detections from the same local trail segment, successive contacts were required to be separated by a minimum longitudinal distance of 3cm.

All agents advanced at constant forward velocity while performing lateral casting movements. Bilateral odor sampling was implemented by evaluating odor concentration at lateral offsets corresponding to the separation of the nostrils. Unless otherwise noted, simulations used a forward velocity of 8 cm/s, inhalation interval of 1/10 s, and bilateral nostril separation of 0.25 cm. Differences between agents arose entirely from how odor encounters were integrated to estimate the continuation of the trail and the uncertainty associated with that estimate.

### Agents with no or limited memory

For a reactive model, the agent ignores all past crossings and predictions, executing only sniff-by-sniff reversals upon loss of odor or high lateral error, mimicking a pure reactive implementation with no memory of the odor trail or concentration. For the limited memory agent, we implemented the model from (Khan et al., 2012) with the exact parameters from their supplementary methods. Briefly the agent advances along the trail at a fixed forward velocity, performing lateral oscillations by alternating the sign of an imposed “casting” acceleration. At each time-step, after an “inhalation”, the agent’s position was updated using: *x*_*t*+1_ = *x*_*t*_ + *v*_*x*_ Δ*t* and *y*_*t*+1_ = *y*_*t*_ + *v*_*y*_ Δ*t* + 1/2 *a*_*y*_ Δ*t*^2^. A reversal of the casting acceleration is either because the odor gradient is greater than the prescribed threshold or the concentration detected drops below the assumed threshold or because the lateral deviation from the current trail estimate surpasses the current uncertainty bound. The agent maintains a running window of the most recent (up to the past 5) trail crossings and maintains an internal estimate of the trail slope.

### Gaussian Process Regression (GPR) agent

The Gaussian Process Regression (GPR) agent treats the trail as a latent spatial function to infer from recent odor trail contacts. Once at least two contacts are obtained, the agent fits a GPR model to remember crossing points (*x*_*k*_, *y*_*k*_, *t*_*k*_), using (*x*_*k*_) as inputs and lateral trail position (*y*_*k*_) as outputs. In pseudocode, the GPR prediction and posterior uncertainty at position *x* was given by: [*ẑ*(*x*), *σ̂*(*x*)] = *GP*. *predict*(*x*). The error estimate of the trail is defined by the GPR prediction ± k posterior standard deviations with k being 2. Briefly, the model treats the trail as a spatially continuous function inferred from discrete odor–trail contact points. These observations are used to condition a Gaussian process framework with a Matérn kernel (smoothness parameter *v* = 1.5, length scale optimized in the interval 5-500 mm). This kernel encodes the assumption that the trail varies smoothly over space while allowing realistic curvature and local deviations. To capture the temporal structure of sampling, recent contacts are weighted more strongly than earlier ones, enabling the estimate to adapt to changes in the trail. The choice of the kernel and the exact parameters are not critical since the model is a phenomenological one designed to recapitulate behavioral data from mice. At every trail contact the GPR was re-fit with the data points weighted by recency given by *w*_*i*_ = exp(−*λ*(*t*_*last*_ − *t*_*i*_)). In addition, since the standard implementation of the scikit-learn GPR does not support weighting samples, an exponential weight of trail contacts is created by replicating each crossing in the training set proportional to *w*_*i*_. Using such an approach, more recent trail contacts are weighted more than temporally older ones. Each agent simulation was performed in batches of 30 independent replicates per condition (which included trail statistics and the four recency weights). For testing the effects of different recency weights, an additional batch of 50 simulations were performed. Each simulation was initialized using a random seed. A memory of 20 most recent detections is used by the GPR agent. All measurements were performed after memory of the agent was saturated with 20 trail contacts so that the internal estimate obtained by the GP fit had stabilized.

The implementation uses the GPR class from Scikit-learn v1.3 (Pedregosa et al., 2011).

### Generalized Worm-Like Chain estimation of the trail (GWLC) agent

The GWLC agent extends the sector-search framework by Reddy et al. (Reddy et al., 2022b) generalizing trail memory and estimate from the two most recent contacts to an arbitrary number of encounters. From these contacts, the agent analytically estimates both the most likely local trail heading and the associated uncertainty, which together defined the orientation and aperture of a sector within which the subsequent search was constrained. As for the GPR agent, the trail was only partially available to the animal, through intermittent odor detections during locomotion. The agent therefore relied on a finite recent history of trail contacts,(*x*_*i*_, *y*_*i*_), retained within a sliding temporal memory window (20 s in the simulations shown). The GWLC model assumes that the trail can be described as a continuous curve whose local heading changes smoothly along space. In particular, the model assumes that curvature fluctuations are Gaussian, have a finite variance, and remain correlated over a characteristic spatial length scale. As a consequence, nearby changes in trail heading are statistically coupled, whereas heading changes over widely separated distances become progressively less correlated. For each segment between successive contact points separated by a distance, these assumptions lead to a Gaussian quadratic form relating the endpoint slopes. The segment-level precision matrix is: *H*_*seg*_ = *α*[[*b*_1_, *b*_2_], [*b*_2_, *b*_1_]] where the coefficients *b*_1_, *b*_2_, and *α* depend on the assumed curvature variance and correlation length of the trail. Segment contributions from all remembered contacts were assembled into a global tridiagonal system spanning the inferred heading variables, and intermediate headings were analytically marginalized by a Schur complement reduction, yielding an effective posterior distribution over the most recent heading estimate. The inferred trail slope and its uncertainty were then given 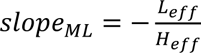 and 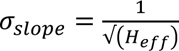, respectively, where *H_eff_* and *L_eff_* denote the marginal precision and linear terms after elimination of earlier variables. The agent advanced at constant forward velocity while performing lateral casting movements. Upon odor contact, the current location was incorporated into memory, and the inferred continuation of the trail was updated. In the absence of further contact, the agent continued to search around the predicted continuation of the trail, carrying forward both an estimate of its expected direction and the uncertainty associated with that estimate. The uncertainty envelope expanded progressively with distance from the most recent contact, and casting direction reversed when the trajectory crossed the inferred bounds surrounding the predicted trail path. In this way, the agent behaved not as a reflexive gradient follower, but as an observer maintaining and continually updating a probabilistic expectation of where the trail ought to lie. A minimum number of contacts was required before GWLC inference became active; prior to this, the agent advanced primarily in the forward direction while GWLC parameters governing curvature variance and correlation length were selected to produce trail statistics comparable to those of the generated environments.

### Extended Kalman Filter (EKF) trail-tracking agent

To estimate the trail heading and the associated uncertainty, the Extended Kalman Filter (EKF) agent samples odor concentration at every sniff through its right and left nostrils. These bilateral, noisy measurements are used to recursively update a probabilistic estimate of the local trail geometry, represented by a latent state vector containing the inferred lateral trail position, local slope, and curvature. The latent trail state is represented as *x*_*t*_ = (*y*_*t*_, *m*_*t*_, *κ*_*t*_), where *y*_*t* denotes the inferred lateral trail position, *m*_*t*_ the local slope, and *κ*_*t*_ the local curvature. State evolution along the forward direction is modeled as a correlated stochastic process with finite spatial correlation length *λ*, the transition model encoded a local second-order expansion of the trail shape, with curvature evolving as an Ornstein–Uhlenbeck process. Between updates in measurement, the state estimate and covariance are propagated forward according to: *x*_*t*+1|*t*_ = *F*(*Δx*)*x*_*t*|*t*_ and *P*_*t*+1|*t*_ = *F*(*Δx*)*P*_*t*|*t*_*F*(*Δx*)^*T*^ + *Q*(*Δx*), where *F*(*Δx*) describes deterministic forward propagation over displacement *Δx*, and *Q*(*Δx*) is the process-noise covariance, computed from the assumed curvature covariance kernel. In this way, uncertainty about trail position increases progressively with distance from recent sensory evidence. Sensory observations are generated through bilateral odor sampling at positions corresponding to the separation of the nostrils. For a given EKF state estimate, the predicted trail position is obtained from the local quadratic expansion 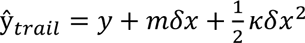, where *δx* denotes the forward displacement of each nostril relative to the agent center. Odor at each nostril is computed from the odor concentration profile defined in the trail generation model. This formulation gave a nonlinear observation model *z*_*t* = ℎ(*x*_*t*_) + *ε*_*t*_, where *z*_*t*_ = (*C*_*L*_, *C*_*R*_) contains the left and right odor measurements,ℎ(*x*_*t*_) predicts the corresponding bilateral concentrations from the current state estimate, and *ε*_*t*_ denotes sensory noise. As the left and right nostrils sample the concentration field at different lateral positions relative to the predicted trail centerline, their difference in concentration provides information about the lateral offset of the trail. Across successive sniffs, these bilateral measurements, combined with the dynamical prior, also constrain the inferred local slope and curvature. Nonlinear measurement updates were computed using a local linearization of the observation model. Specifically, the Jacobian was used to compute the Kalman gain 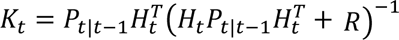 and to update both the state estimate and covariance. Thus, each bilateral odor sample acted not simply as a local cue, but as evidence constraining an ongoing probabilistic estimate of the unknown trail position. The agent advanced at constant forward velocity while performing lateral casting movements. Following odor encounters, the lateral position estimate was anchored toward the detected trail location, and the associated positional uncertainty was reduced locally while preserving the ongoing dynamical estimate of slope and curvature. In the absence of further contact, the agent continued to search around the predicted continuation of the trail. The extent of casting was determined by the inferred uncertainty envelope surrounding the estimated trail position, and casting direction reversed whenever the trajectory crossed the predicted uncertainty bounds. A minimum number of odor encounters was required before uncertainty-guided casting became active. Prior to this, the agent advanced primarily in the forward direction. Odor encounters themselves were probabilistic, with detection likelihood determined by local odor strength. Trail statistics were defined by the spatial correlation length and curvature variance parameters of the EKF prior. Unlike the GPR or the GWLC agent, EKF itself was recursive, such that past observations influenced future estimates only through the evolving latent state and covariance matrix without an explicit memory of the previous detections.

### Glomerular imaging

To optically access the glomeruli, a craniotomy was performed above the olfactory bulb of mice expressing jGCaMP8m (JAX #037718) in olfactory sensory neurons using *OMP-Cre* mice. Mice were anesthetized with an intraperitoneal injection of ketamine and xylazine (100 and 10 mg/kg, respectively), and petroleum jelly was used on the eyes to keep them hydrated. The temperature of the body was maintained at 37 °C using a heating pad. The scalp was shaved and then opened with a scalpel blade. After thorough cleaning and drying, a craniotomy was performed over the OBs using a 3 mm diameter biopsy punch (Integra Miltex). The surface of the brain was cleared of debris. The surface of the brain was kept moist with artificial cerebrospinal fluid containing in mM (125 NaCl, 5 KCl, 10 Glucose, 10 HEPES, 2 CaCl2, and 2 MgSO4 [pH 7.4]) and Gelfoam (Patterson Veterinary). A coverslip pair was made by gluing a 4mm No. 1 coverslip with a 3mm No. 1 glass coverslip (Warner) with optical adhesive (Norland Optical Adhesive 61). The coverslips were adhered to the edges of the cranial cavity in the skull with Vetbond (3 M). For obtaining respiration measurements, we also inserted thermistors in the same surgery as described above. The posterior portion of the exposed skull was gently scratched with a blade, and a custom, 3D printed half head bar was attached using C&B-Metabond dental cement (Parkell, Inc.). Buprenorphine extended-release (3.25 mg/kg) (Ethiqa XR) and Carprofen (5 mg/kg) were administered subcutaneously for analgesia at the end of the surgery.

### Widefield calcium imaging

To obtain neurophysiology with cellular resolution we performed widefield calcium imaging using head mounted miniature microscopes. For glomerular recordings over the olfactory bulb (OB), a UCLA v4 miniscope (board v4.3, OEPS) was assembled, and resolution targets were imaged to ensure that the Field of view (FOV) was clean and the electronic focus worked. The thermistor cables were wrapped around the coax cable used for the miniscope and the coax cable was routed through a custom-built commutator which was used to ensure the cable was not twisting especially in the early sessions. For recordings in the AON, an Inscopix (nVista3, Inscopix) was used, and the plane was adjusted to obtain the most neurons.

### GRIN lens implantation

For imaging neurons in the AON, 400 nl of AAV9-syn-jGCaMP8m-WPRE (Addgene, 162375-AAV9) was injected unilaterally into the C57BL/6J mice, into the AON (AP-2.9, ML-1.2, DV- 2.7 from the brain surface. After the virus was injected using a pulled glass pipette attached to a nanoinjector (MO-10, Narishige), either a 25-gauge blunt needle (McMaster-Carr 6710A33, for 0.6mm lens implants) or a bone chisel with a 1mm cutting width (Fine Science Tools #10096-16, for 1mm lens implants) was slowly inserted (1 mm per 5 min) into the brain targeting the same AP and ML coordinates and DV-2.5 to create a tract. The needle then was withdrawn slowly, and a GRIN lens (either Inscopix, 0.6 × 4.3 mm or Inscopix 1mmX 4mm) was inserted slowly (1 mm per 10 min) into the tract formed by the needle and targeted at 100 µm below the end of the needle tract. The lens assembly with the baseplate was secured on the skull with C&B-Metabond dental cement (Parkell, Inc.), and a custom, 3d printed head bar was attached at the base of the lens assembly with dental cement. This allowed brief restriction to attach the microscope to the animal’s head. Buprenorphine extended-release (3.25 mg/kg) (Ethiqa XR) and Carprofen (5 mg/kg) were administered subcutaneously for analgesia at the end of the surgery. The positions of all implanted GRIN lenses were assessed post hoc through histology and only the imaging data from correctly targeted lens were used for further analysis. Mice were housed individually after surgery and were given 3–4 weeks to allow for the expression of GCaMP and clearing of the imaging window.

### Image processing and calcium signal extraction

Recordings from the olfactory bulb were acquired using a UCLA v4 miniscope system at 30 Hz. Imaging data were synchronized with behavioral cameras via TTL triggers to a data acquisition system, and movies were recorded using Bonsai. Movies were motion corrected, spatially filtered, and calcium sources were extracted using the CNMF-E algorithm implemented in CaImAn. Recordings from the anterior olfactory nucleus (AON) were acquired using the Inscopix nVista3 system at 30 Hz and similarly synchronized with behavioral cameras via TTL triggers. Raw imaging files were spatially down sampled (4x), motion corrected and spatially filtered using Inscopix Data Processing Software (IDPS) before being exported as TIFF stacks. Calcium sources were then extracted using the CNMF-E algorithm in CaImAn. Extracted components were manually curated based on spatial footprint, temporal trace quality, signal-to-noise ratio, peak-to-noise ratio, motion artifacts, decay kinetics, and anatomical plausibility. Components with abnormal morphology, poor signal quality, or clear motion-related artifacts were excluded. For neuronal datasets, elongated or thin components inconsistent with expected cell morphology were removed.

### Binary decoding analyses

To test whether population activity at a given sniff could predict whether the animal was near or far from the trail, population activity vectors, each dimension corresponding to one glomerulus/ROI, were used to train logistic regression classifiers on sniff-by-sniff data. Sniffs were labeled according to whether the animal was within or beyond 1.5 cm of the trail, and decoder performance was evaluated using stratified 5-fold cross-validation with out-of-fold predictions, quantified by balanced accuracy and ROC AUC. To examine how information scaled with population size, decoding was repeated across 100 subsets for randomly chosen subsets of ROIs without replacement, providing an estimate of variability due to ROI choice. Chance performance was estimated by repeating the same procedure after randomly permuting class labels. To decode the position of the snout relative to the trail, sniffs were instead labeled according to the signed lateral deviation of the snout from the trail centerline. Sniffs within ±1 cm of the trail center were excluded and left and right samples were matched within 0.5 cm-wide bins of absolute snout deviation prior to decoding to control for differences in distance from the trail.

### Single-unit and population activity analysis

All analyses of extracted calcium traces were performed using custom Python scripts. Calcium traces were organized by animal, session, imaging field, behavioral epoch, and trial identity. For turn-related analyses, activity from each extracted unit was aligned to the onset of behavioral turns, and peri-event calcium activity was quantified within defined time windows surrounding each turn. Baseline activity was calculated from the pre-event period and turn-related responses were assessed by comparing peri-turn activity to this baseline period. For straight-section analyses, trials were grouped according to the RMS of the preceding trail, a measure of its recent tortuosity. Straight segments following trails with different RMS values were used to assess whether neural activity reflected recent trail statistics. Calcium activity during these straight sections was extracted and compared across preceding-trial RMS-defined conditions.

### Data analysis and Statistics

Markerless pose estimation was performed using DeepLabCut v2.2.0 (Mathis et al., 2018; Nath et al., 2019) to obtain body parts of interest, viz. the snout (the tip of the nose), the ears, the shoulder and the base of the tail. For the view from above, 950 frames taken from 36 videos for 350,000 training iterations with ResNet-50v1 and the train and test errors were 6.69,7.85 pixels respectively with a 0.1 p-cutoff (2048 x 1400 image size). For the view from the side, 380 frames taken from 8 videos for 150,000 training iterations with Restnet-101 and the train and test errors were 5.91 and 20.03 with a 0.1 p cutoff (2048 x 760 image size). The trail was also detected using DeepLabCut to identify the edges of the trail, and other features such as rewards. 3000 frames taken from 40 videos were used with ResNet-101 for 450,000 training iterations and the train and test errors were 21.63 and 24.53 with a p-cutoff of 0.1 (1504 x 608 image size). Sessions were reconstructed by “rolling out” using an odometer that used a rotary absolute shaft encoder (US digital MA3) connected through a wheel placed on the paper. The encoder was read out using the same microcontroller controlling the DC motor and was interfaced with the acquisition computer using serial communication. For estimating the elevation of the snout, another network was trained to detect the limbs of the animal. For this, 670 frames taken from 17 videos for 200,000 training iterations with RestNet-50v1 and the train and test errors were 2.6, 12.36 pixels with a 0.1 p-cutoff (2048 x 760 image size). The training was done using an NVIDIA Titan V or a 2080Ti GPU. After obtaining the x, y coordinates of their hands and feet, a straight line was approximated to obtain the fore-hind paw plane using RANSAC (RANdom SAmple Consensus) robust linear regression implementation to reduce the sensitivity to the outlier points. The orientation (slope and intercept) of this fitted line provided an estimate of the mouse’s base of support at that instant. Finally, the vertical intersection between the x coordinate of the snout and the fitted line was computed and plotted back on the frame for manual verification.

For estimation of respiration, signals were first filtered using a Butterworth filter to remove high frequency noise, following which a Hilbert transform was applied to obtain the envelope. Peaks and troughs were then detected using the *find_peaks* function in SciPy (Virtanen et al., 2020).

No statistical method was used to predetermine sample size. Statistical details including tests and p-values can be found in the figure legends and results section. Numbers of animals are reported in the text or the figure legend. In the text, we report mean ± s.e.m. (standard error of the mean) for datasets where means were compared. Jackknife resampling was used to estimate standard errors for percent tracking, with each animal left out in turn. All comparisons were performed at the animal level using summary statistics averaged across trials within each animal. When a two-sample Kolmogorov-Smirnov test was used to assess group differences, significance was assessed at the animal-level and p-values were obtained by permutation (1,000 permutations) rather than from the asymptotic distribution. The KS statistic D and associated permutation-based p-values are reported throughout.

## Acknowledgements

We would like to thank Ed Soucy and Brett Graham for their technical support. We thank Gautam Reddy and Jacob Zavatone-Veth for helpful discussions. We also thank Andrew Meng, Vikrant Kapoor, Jonah Pearl, and members of the Murthy lab for valuable feedback. We would also like to thank the Harvard Center for Biological Imaging (RRID:SCR_018673) for infrastructure and support. This work was supported by Harvard University, by DFG grant MA 6176/1-1 (A.M.), and by a Marie Curie Fellowship PIOF-GA-2013-622943 (A.M.). This research was supported in part by the Gordon and Betty Moore Foundation Grant No. 2919.02 and NSF grant PHY-2309135 to the Kavli Institute for Theoretical Physics (KITP). Research in V.N.M.’s lab was supported by grants from the NIH (RF1NS128865) and NTT Research (A47994). Work by N.R. and M.V. was supported by NIH (RF1NS128865).

## Author Contributions

S.J., A.M., M.V. and V.N.M. conceived the project. A.M. designed and built the initial experimental setup and piloted experiments with M.W.M. S.J. refined and adapted the behavioral rig, collected and analyzed the data reported in this paper, with input from N.R., A.M., M.V., and V.N.M. S.J. performed the imaging experiments, using the GRIN lens approach piloted by M.M.R., who also provided surgical support. S.J. and N.R. implemented the model with inputs from M.V. and V.N.M. S.J. and V.N.M. wrote the manuscript with edits and inputs from all the authors.

## Supplementary Figures

**Figure S1:**
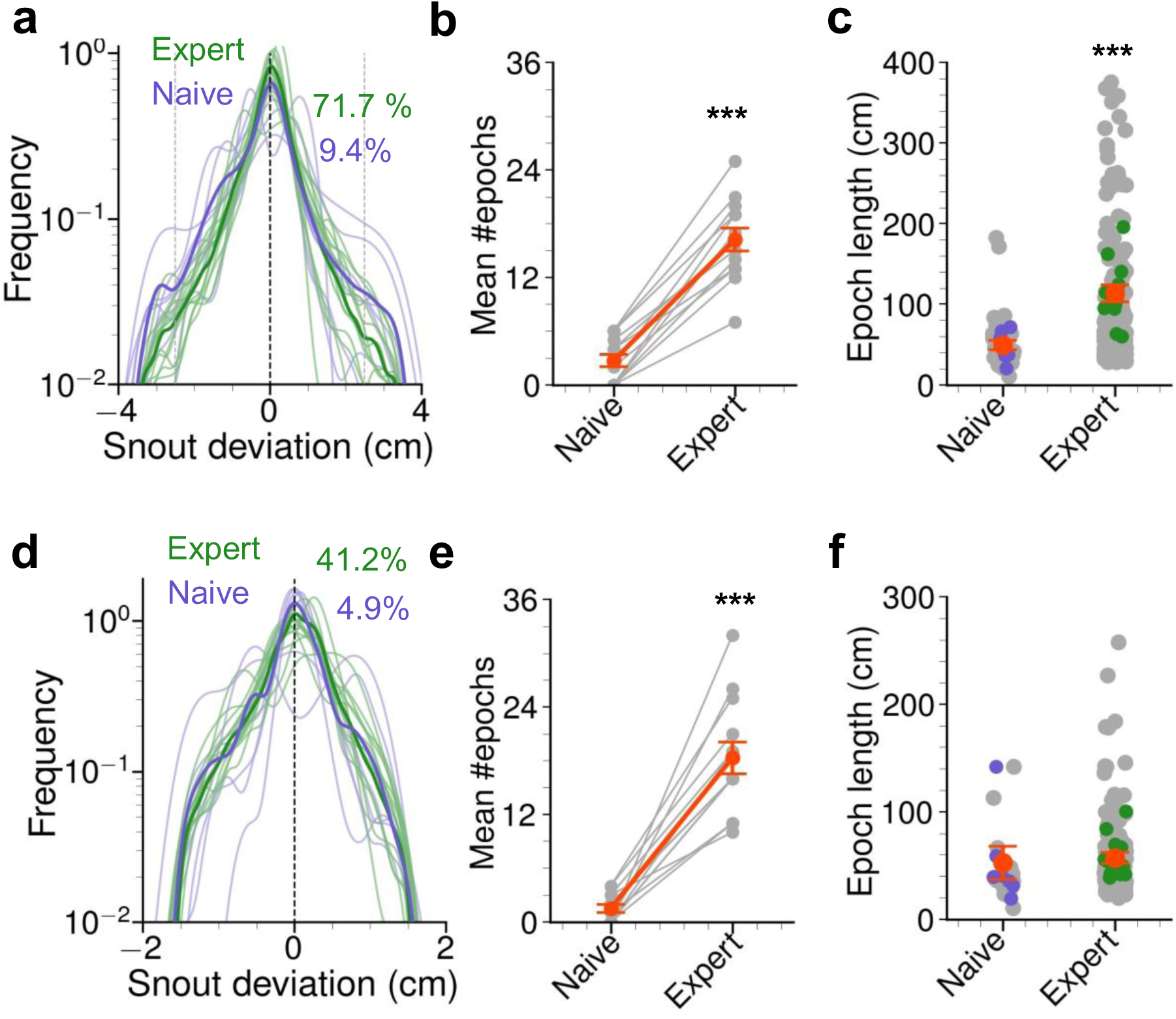
Trail tracking differences across learning are not influenced by selection criteria. Parameters of following a trail with two selection criteria that are different from that shown in Figure 1d. 3.5 cm for 2.5 seconds in panels a-c, and 1.5cm for 2.5 seconds in panels d-f. Each line represents a separate animal (**a)** The same distribution shown in Figure 1c limited to 3.5 cm on either side of the trail. Distributions of the deviations of the snout from the trail. Numbers indicate percentage tracking. (**b)** Mean number of epochs. Gray points represent 13 individual animals and red is average. **(c)** Mean length of epochs. The colored dots represent individual animals. **(d)** The same distribution shown in Figure 1c limited to 1.5cm on either side of the trail. Numbers indicate percentage tracking. **(e)** Mean number of epochs. **(f)** Mean length of epochs. These are the same animals shown in panel d-f of Figure 1 (13 animals).

**Figure S2:**
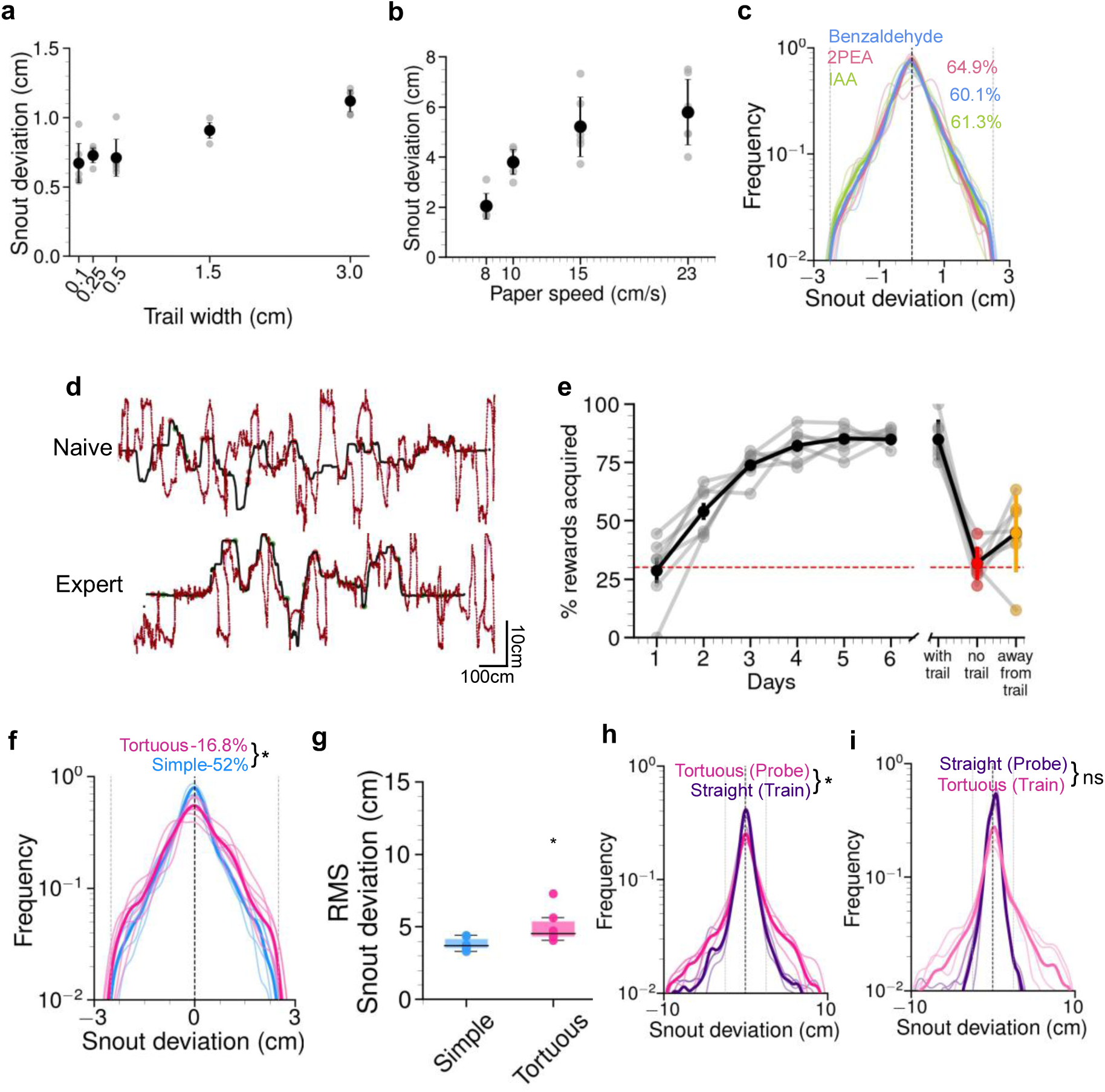
Factors that influence trail tracking. **(a)** Average deviation from the trail from different sessions with varying widths of the trail displayed as mean ± s.e.m (6 animals). Each gray dot is a separate animal. **(b)** Average deviation from the trail as a function of the speed of the paper displayed as mean ± s.e.m. Each gray dot is a separate animal. **(c)** Distributions of the deviations of the snout from the trail from mice that were presented with different odorants. The numbers indicate percentage tracking. **(d)** An example entire session from a mouse following a trail on day 1 (naïve) and on day 6 (expert). **(e)** Percentage of rewards consumed while following odor trails over days shown as average ± s.e.m.(7 animals) **(f)** Distributions of the snout deviations from the trail of mice that were following simple or tortuous trails. **(g)** Root mean square deviations from the trail from animals tracking either a simple straight trail or a tortuous trail.(ns: not significant, p =0.17, Mann-Whitney U test, 5 mice). (**h, i**) Distributions of snout deviation from 4 mice that were trained on only straight trails and tortuous trails were used as a probe session (**h**) (D=0.11,*p<0.05, KS test) or were trained on tortuous trails and were tested on straight trails (**i**) (ns:not significant).

**Figure S3:**
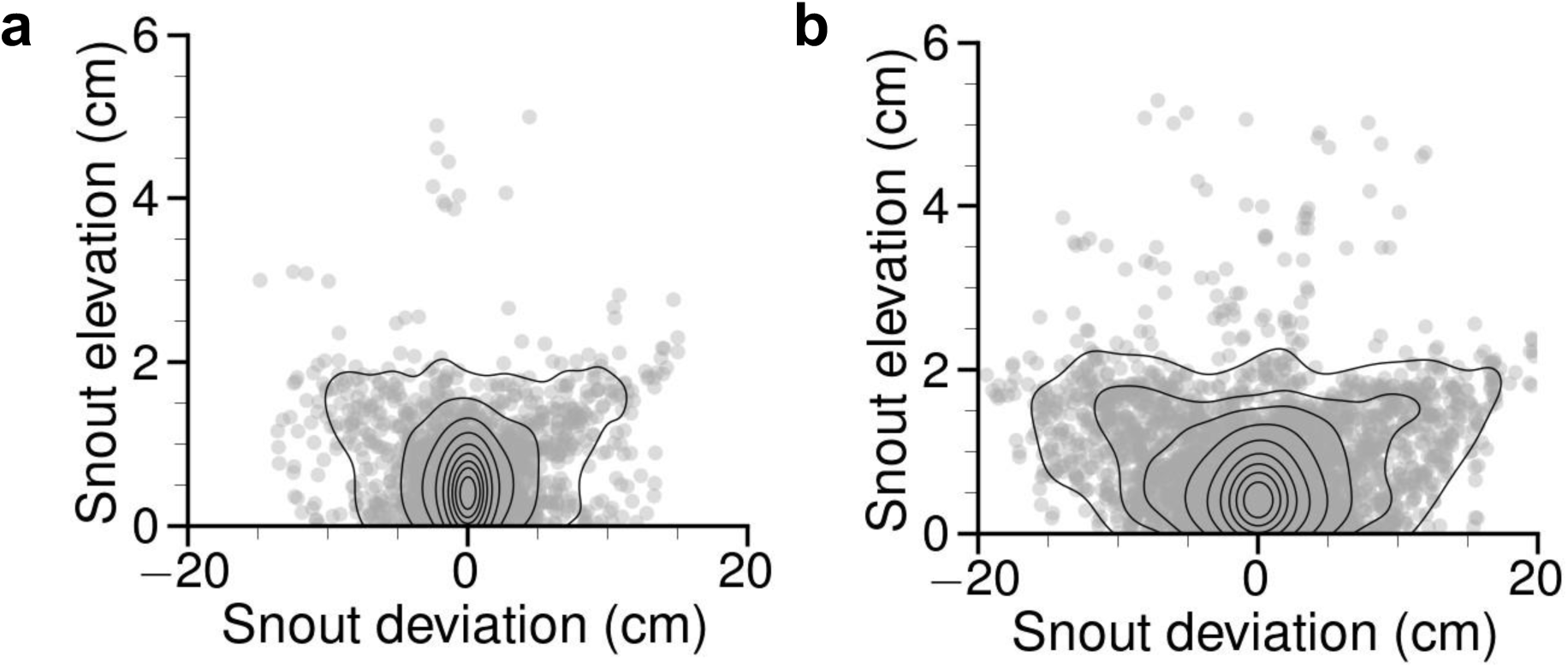
Introduction of the trail odor in the air disrupts trail tracking. Distributions showing snout occupancy in 3d space without **(a)** and with **(b)** the trail odor presented aerially. The distribution is combined from 9 animals.

**Figure S4:**
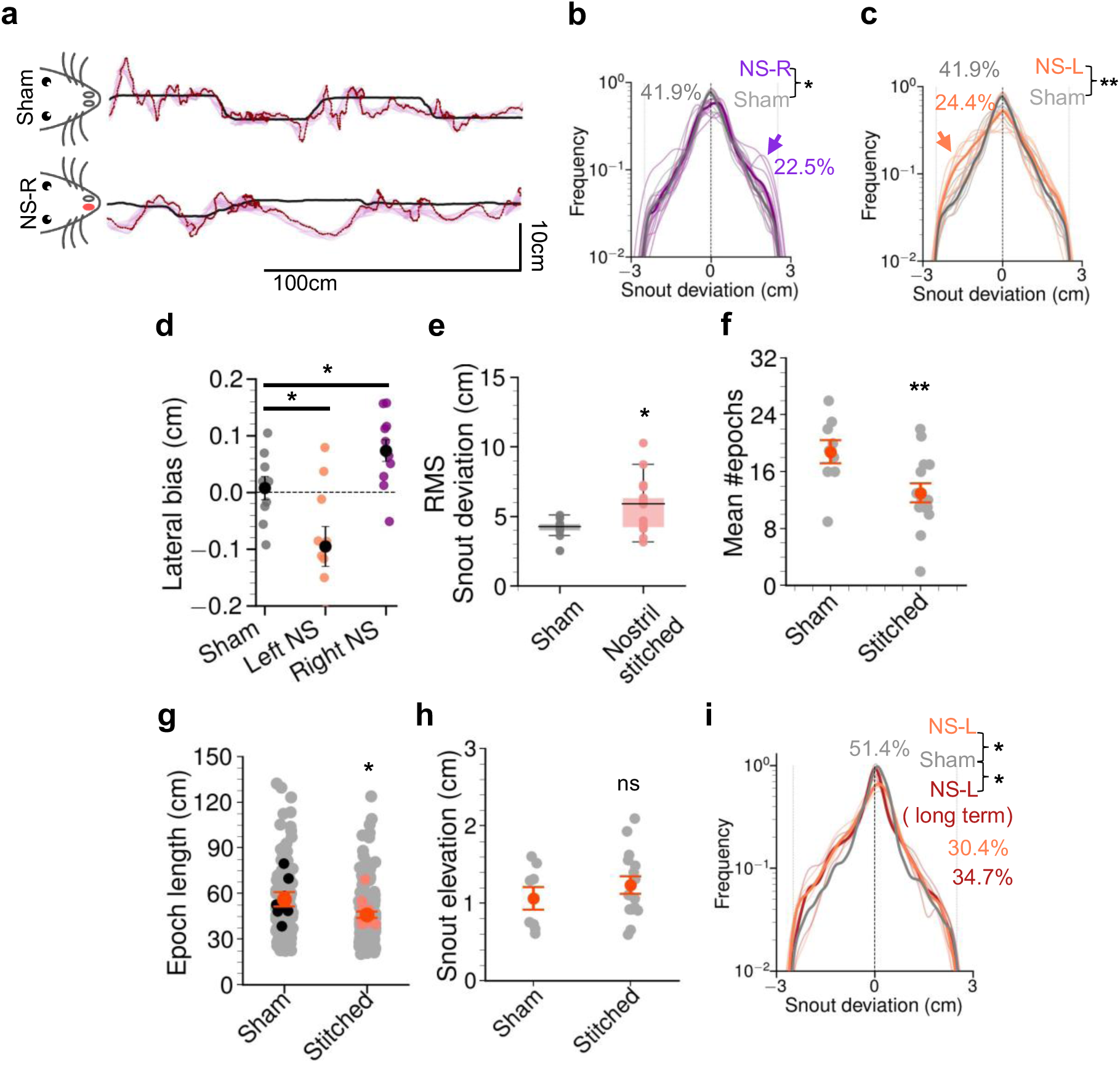
Blocking the nostril disrupts trail tracking. **(a)** Example trajectory from a mouse following a trail after sham stitch (see Methods) or when the right nostril is stitched. **(b** and **c)** Distributions of the deviations of the snout from the trail from mice with a sham stitch and when the right nostril was stitched **(b)** or when the left nostril was stitched **(c)**. The arrow indicates bias toward the closed side. Numbers indicate percentage tracking. Individual lines show each animal. Panel (c) is reproduced from Figure 2c to allow direct comparison between the sides. (**d**) Figure shows lateral bias, which was quantified as the signed lateral offset between the snout and the nearest point on the odor trail during tracking epochs. Positive values indicate rightward offsets relative to the trail and negative values indicate leftward offsets. Individual dots represent unique animals. (**e-h)** Performance metrics of trail tracking for sham and mice with a single nostril stitched. Individual dots in **e**, **f** and **h** represent unique animals as do the colored dots in **g**. **(e)** Root mean square deviation from the trail from either sham or nostril stitched conditions. **(f)** Mean number of epochs. **(g)** Mean length of epochs and **(h)** Elevation of the snout from the paper floor. **(i)** Distributions of the deviations of the snout from the trail from 4 mice immediately after the nostril was stitched (coral) and after 7 days of keeping the nostril stitched (red). (**\***p<0.05,**\*\***p<0.01, ns: not significant, Wilcoxon rank-sum test).

**Figure S5:**
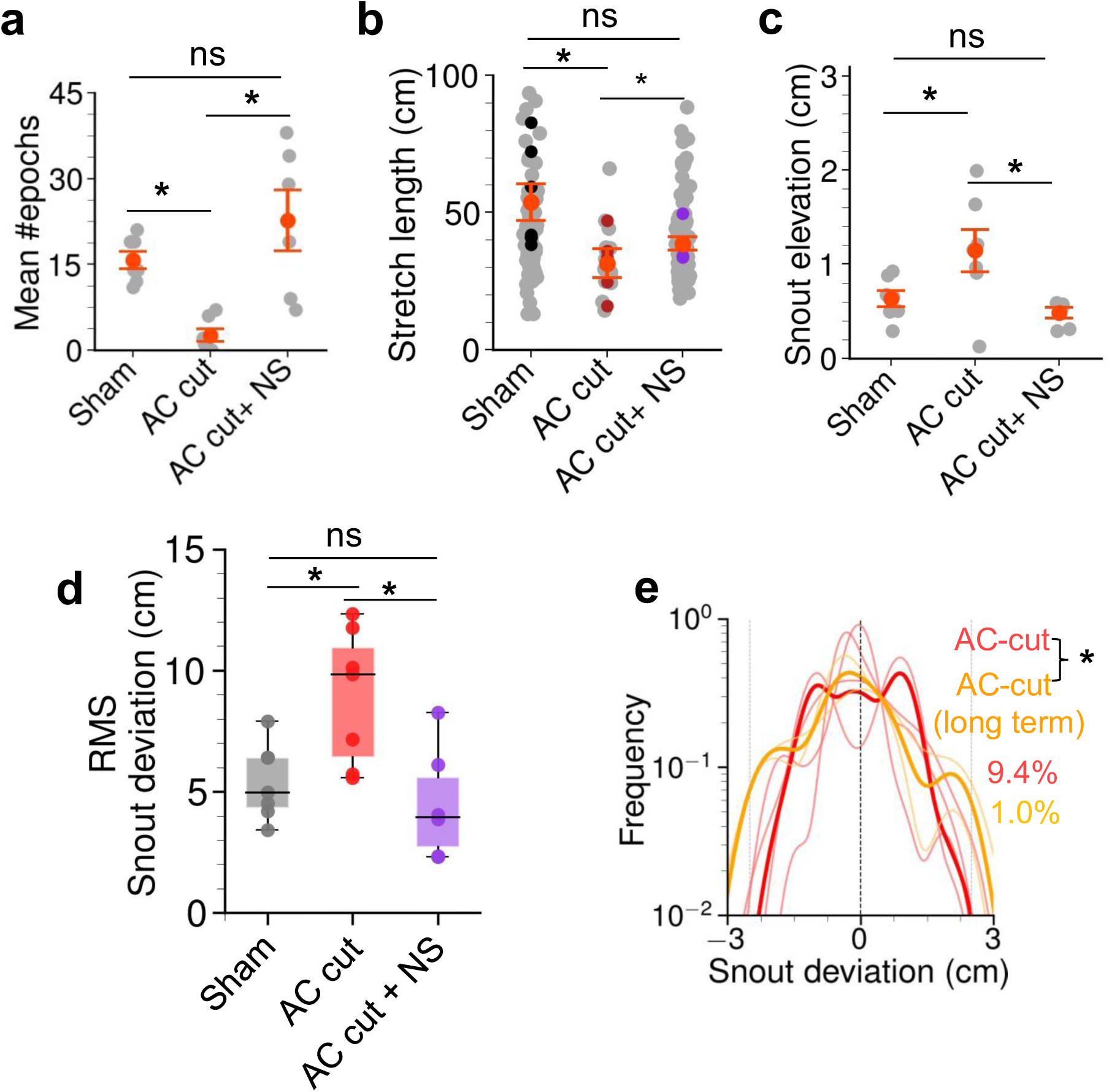
Summary of trail tracking measures after the anterior commissure was severed. (**a-c)** Performance metrics of trail following between sham, mice with anterior commissure severed and with anterior commissure severed but with one of the nostrils stitched. Gray dots in a,c as well as colored dots in b represent unique animals. **(a)** Mean number of epochs. **(b)** Mean length of epochs and **(c)** Elevation of the snout from the paper floor. **(d)** Root mean square deviation from the trail from either sham, animals with the commissure transected or animals with the anterior commissure transected and one of the nostrils stitched. **(e)** Distributions of the deviations of the snout from the trail from mice where the anterior commissure was severed immediately after (red) and after 2 weeks (orange). Lines represent individual mice (**\*\***p<0.01, **\***p<0.05, ns: not significant Wilcoxon rank-sum test). Numbers indicate percentage of time mice spend tracking (as defined in Methods).

**Figure S6:**
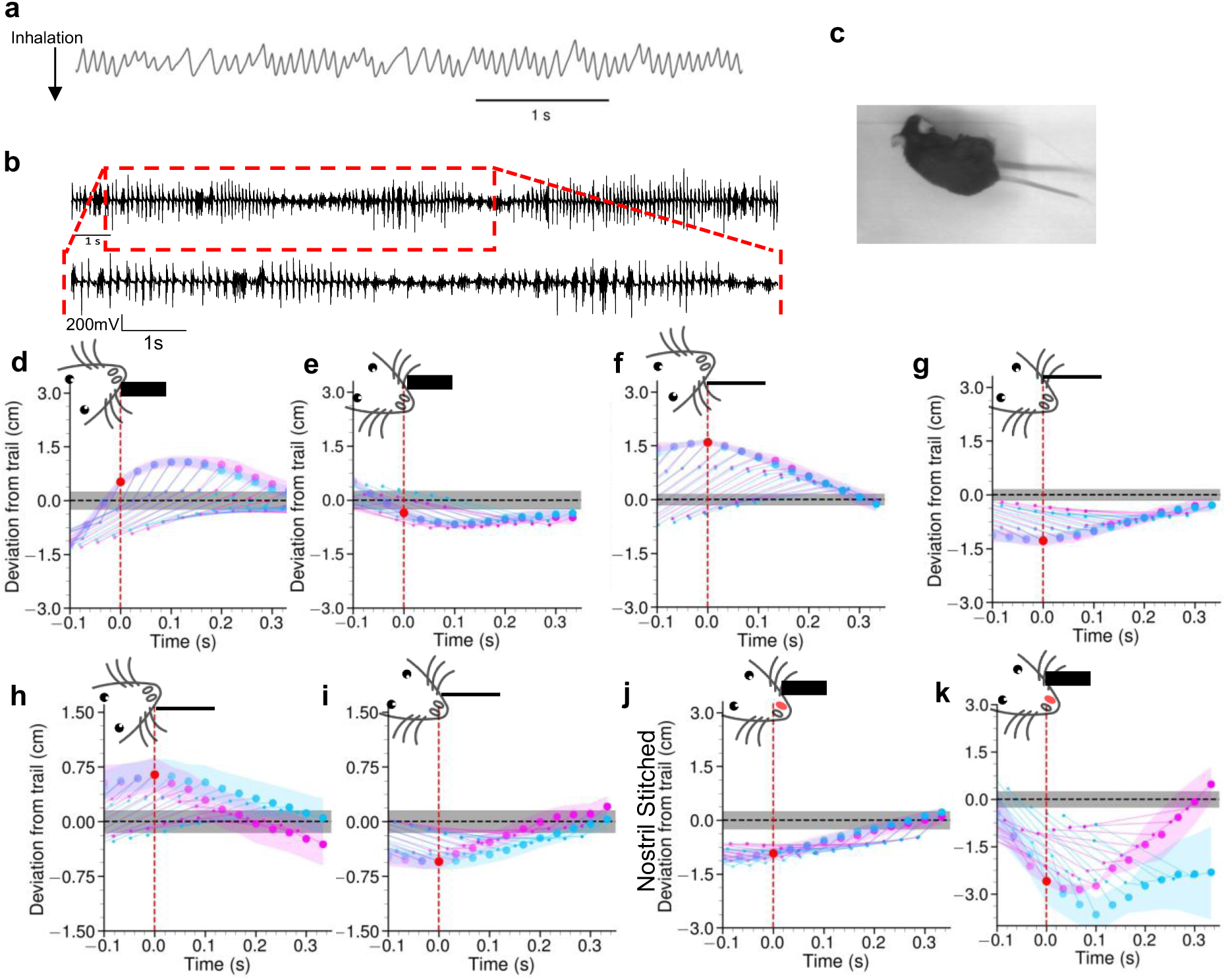
Measuring respiration during trail tracking. **(a)** Example time series from a thermocouple from a mouse following the trail. **(b)** Example EMG trace from a mouse implanted with a wireless EMG implant, with the inset showing a portion of the trace at higher resolution. **(c)** Image showing a mouse implanted with the wireless EMG sensor following a trail. **(d** and **e)** Trajectories showing snout, shoulder and base of the tail, joined by lines. The red dot indicates an inhalation event. Pink traces are from when an inhalation was detected at t = 0, and cyan traces are when there is no inhalation at t = 0. The trail is 0.5cm in width and shown as a gray line. The distance at the inhalation is between 0 and 1cm and is on the trail. (**f** and **g**) Trajectories showing snout, shoulder and base of the tail, joined by lines from a trail whose width is 0.2cm as shown by the gray line. (**h** and **i**) show trajectories when the inhalation event is closer to the narrower trail. (**j** and **k**) show trajectories when the inhalation event is closer or further away from a trail of width 0.5cm in mice where the left nostril is stitched. (at least 25 stretches from 5 mice).

**Figure S7:**
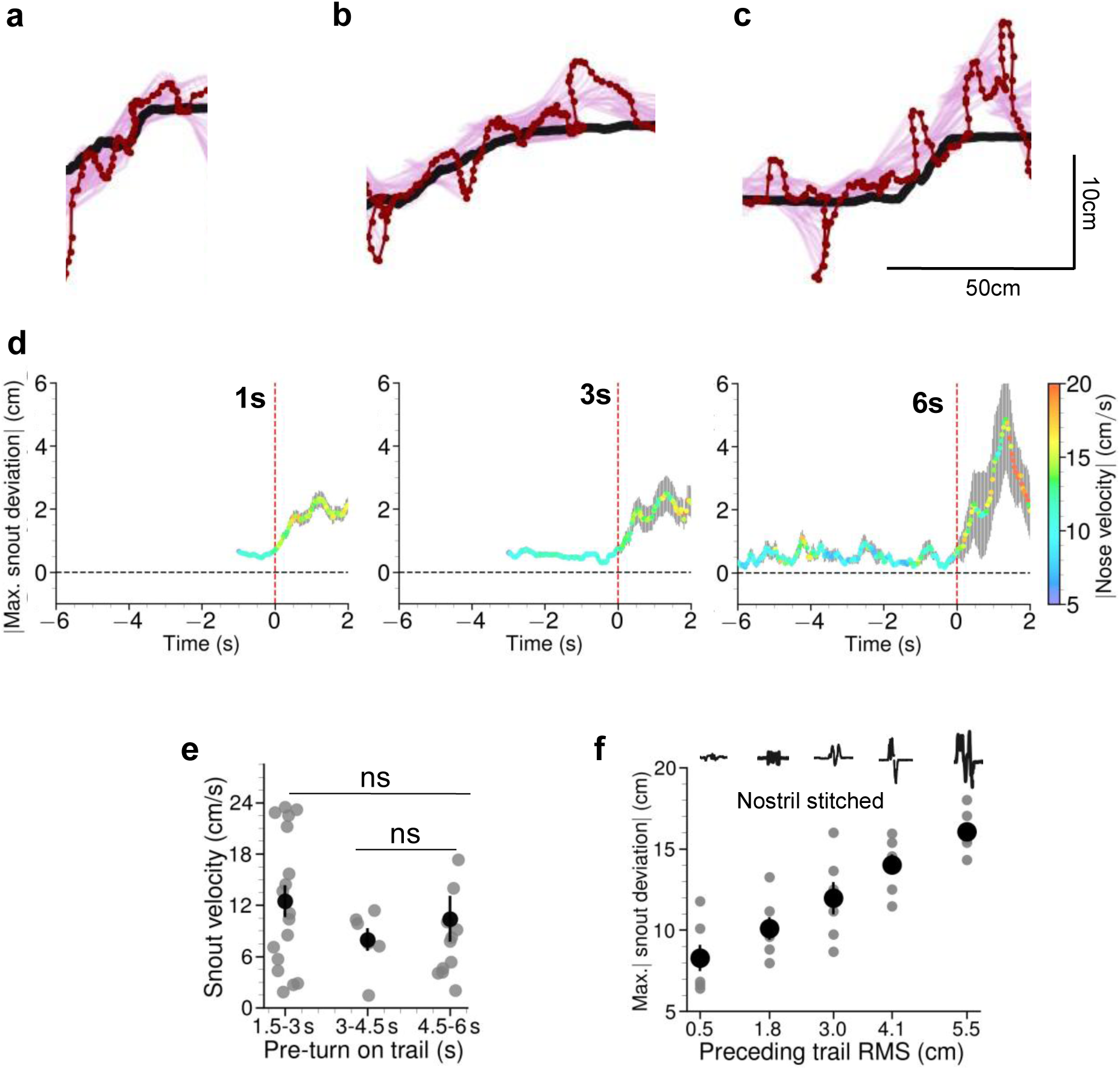
Off times for memory post encountering noisy trails. **(a,b,c)** Example trajectories from a mouse encountering a turn in the trail, after following the trail for 1s **(a)**,3s **(b)** or 6s **(c)**. **(d)** Snout positions over time shown in Figure 4e colored by nose velocity. **(e)** Summary showing the average velocity of the nose at the point when the trail turns, binned into 1.5 second bins. These are not significantly different (p>0.8, Kruskal-Wallis ANOVA with post hoc Mann-Whitney U tests using Bonferroni multiple comparisons correction). **(f)** Maximum snout deviation as a function of preceding trail RMS from 4 animals whose nostril was stitched.

**Figure S8:**
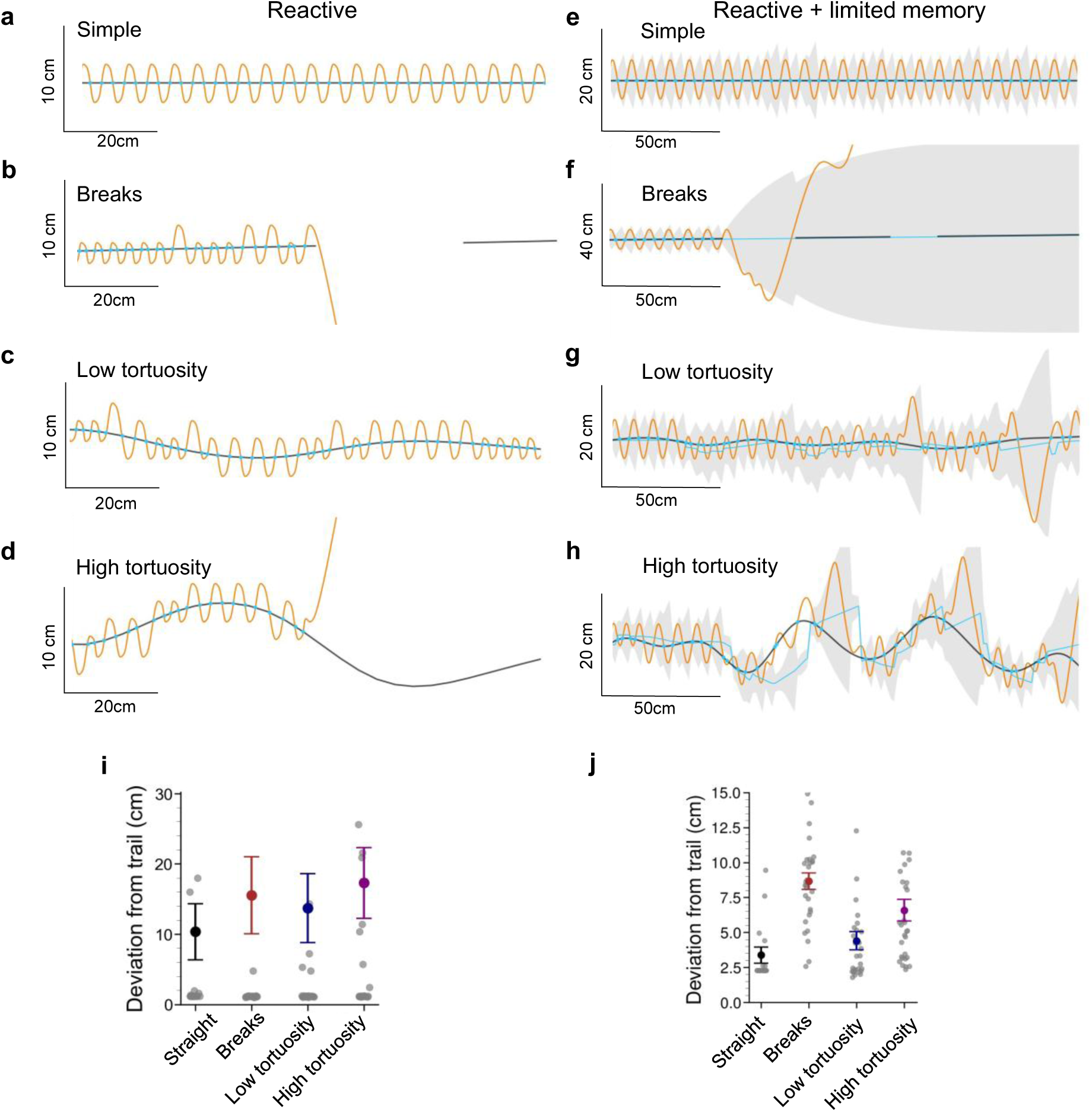
Reactive models and models with limited memory can track trails with limitations. **(a-d)** Example trajectories of a reactive agent with no memory following a trail that is straight (a), simple with breaks (b), of low tortuosity (c) or high tortuosity (d). **(e-h)** Example trajectories of an agent that is reactive but also has up to 5 previous contact points in memory to form an estimate of the trail that was presented a trail that is straight (e), simple with breaks (f), of low tortuosity (g) or high tortuosity (h). **(i)** Snout deviations from 30 simulations across the various trail types shown through a-d. **(j)** Snout deviations from 30 simulations across the various trail types shown through e-h.

**Figure S9:**
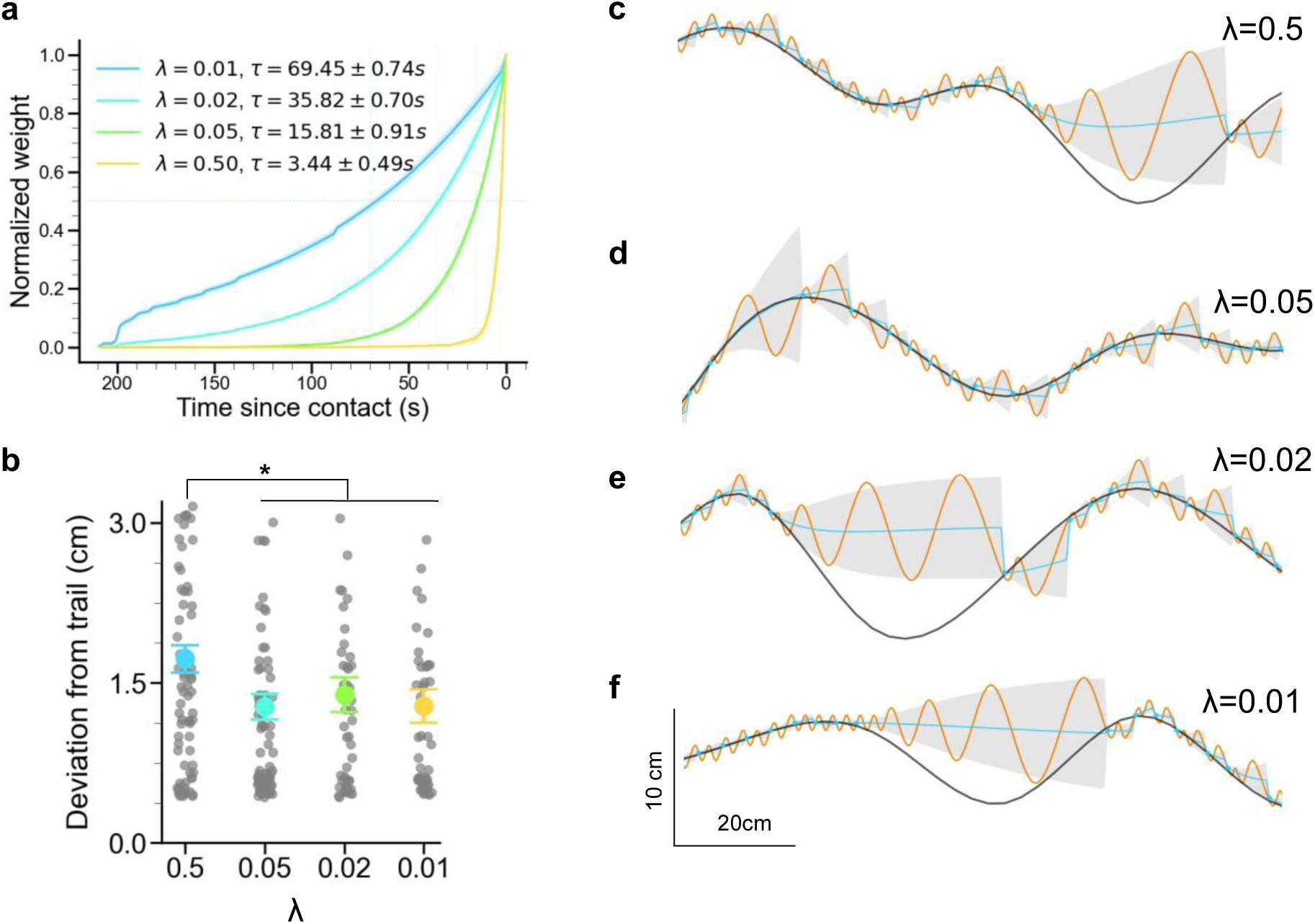
Agents with memory. **(a)** Normalized weight of previous contacts plotted against time since last contact from 30 agent simulations for the same four values of λ shown in (c-f). The most recent contact is on the right. **(b)** Deviations from the trail shown as mean ± s.e.m from at least 80 simulations of agents following a trail of high tortuosity for the same four values of λ. The snout deviation of the agents with a λ value of 0.5 is significantly higher than λ = 0.05,0.02 or 0.01. **\***p<0.05, Kruskal-Wallis ANOVA with post hoc Mann-Whitney U tests using Bonferroni multiple comparisons correction test. **(c-f)** Example trajectories from an agent following a fictive trail of high tortuosity with varying lambdas λ that determine how many past contacts are weighted.

**Figure S10:**
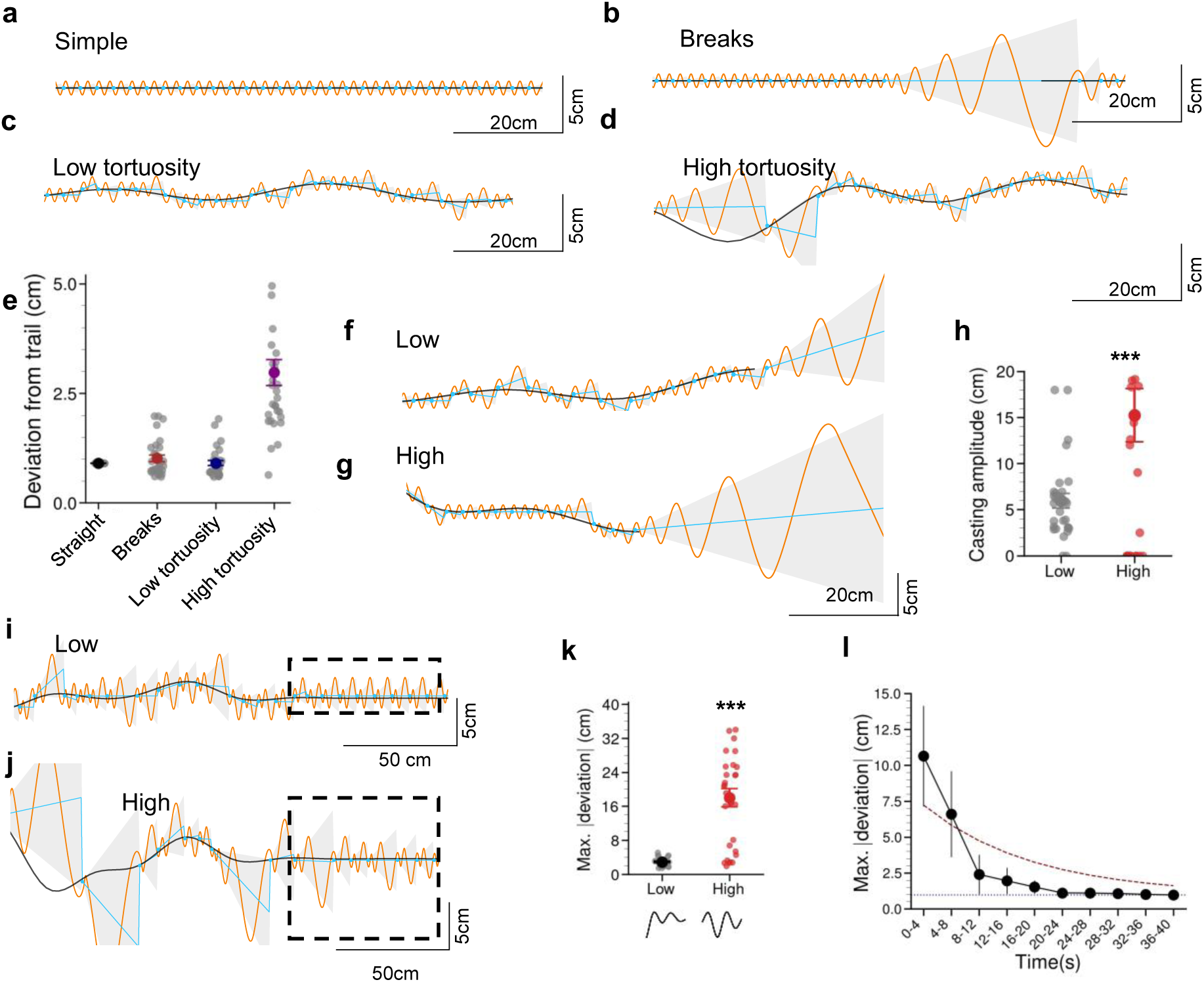
An agent using memory reproduces many facets of tracking behavior. **(a-d)** Example trajectories of an agent using a Generalized Worm-Like Chain (GWLC) to estimate the trail from previous contacts with the trail. The virtual trail is shown in black, the agent’s trajectory in orange, the agent’s trail encounters as sky blue dots, the agent’s heading estimate in a sky blue line and the posterior uncertainty of the trail position in gray cones. The example trajectories are from the agent following **a)** a simple straight line, **b)** a simple trail with breaks, **c)** curves of low tortuosity and **d)** curves of high tortuosity. **(e)** Deviations from the trail as shown by means ± s.e.m. are from 30 simulations, show that the agent is accurate in following trails. **(f** and **g)** Example trajectories of the agent encountering a break while following trails of two types of curvature. **(h)** Casting amplitudes measured from 30 simulations as shown by means ± s.e.m. show that the agent casts more widely when encountering a break following the trail of higher curvature. **\*\*\***p<0.001, Mann-Whitney U test. **(i** and **j)** Example trajectories from an agent following a straight section of a trail after encountering either low tortuosity **(i)** or high tortuosity **(j)**. **(k)** Snout deviations over time from agents following a trail. The deviations are from straight sections of the trail. The deviations are significantly higher when the trail was preceded by the curves of higher tortuosity. **(l)** Maximum deviation of the agent in a straight section of the trail immediately after encountering a trail with a root mean squared deviation of 3cm. The fitting time constant shown is 15.8 seconds. **\*\***p<0.01**, \***p<0.05, Mann-Whitney U test.

**Figure S11:**
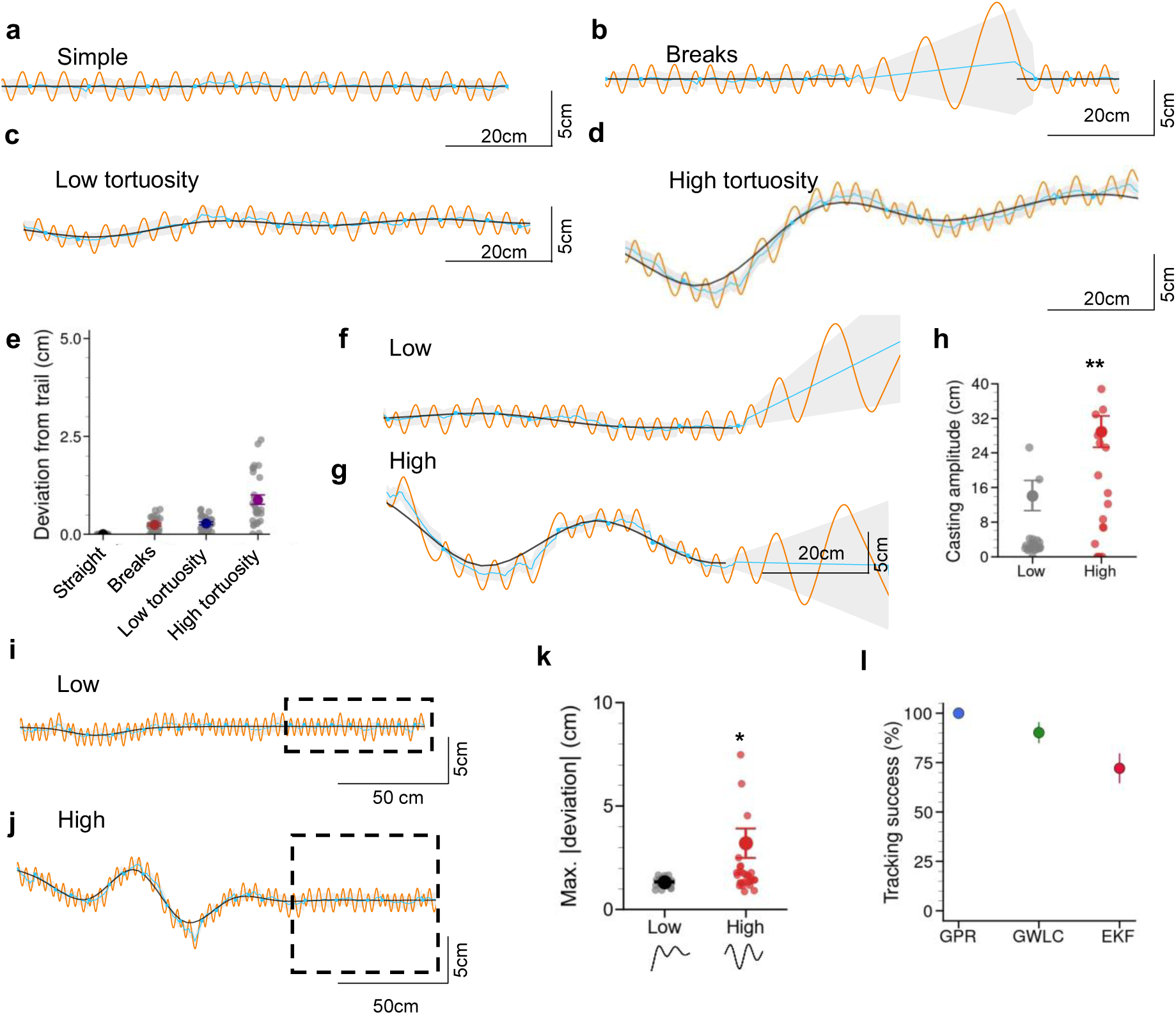
An agent using memory based on an Extended Kalman Filter reproduces some facets of tracking behavior. **(a-d)** Example trajectories of an agent using an Extended Kalman Filter (EKF) to estimate the trail from previous contacts with the trail. The virtual trail is shown in black, the agent’s trajectory in orange, the agent’s trail encounters as sky blue dots, the agent’s heading estimate in a sky blue line and the posterior uncertainty of the trail position in gray cones. The example trajectories are from the agent following **a)** a simple straight line, **b)** a simple trail with breaks, **c)** curves of low tortuosity and **d)** curves of high tortuosity. **(e)** Deviations from the trail as shown by means ± s.e.m. are from 30 simulations, show that the agent is accurate in following trails. **(f** and **g)** Example trajectories of the agent encountering a break while following trails of two types of curvature. **(h)** Casting amplitudes measured from 30 simulations as shown by means ± s.e.m. show that the agent casts more widely when encountering a break following the trail of higher curvature. **\*\*\***p<0.001, Mann-Whitney U test. **(i** and **j)** Example trajectories from an agent following a straight section of a trail after encountering either low tortuosity **(i)** or high tortuosity **(j)**. **(k)** Snout deviations over time from agents following a trail. The deviations are from straight sections of the trail. The deviations are significant when the trail was preceded by the curves of higher tortuosity. **(l)** Tracking success of either the GPR, GWLC or EKF agent defined as not deviating more than 20cm from the trail in the tortuous trail and succeeding to make it to the straight section. **\*\***p<0.01**, \***p<0.05, Mann-Whitney U test.

**Figure S12:**
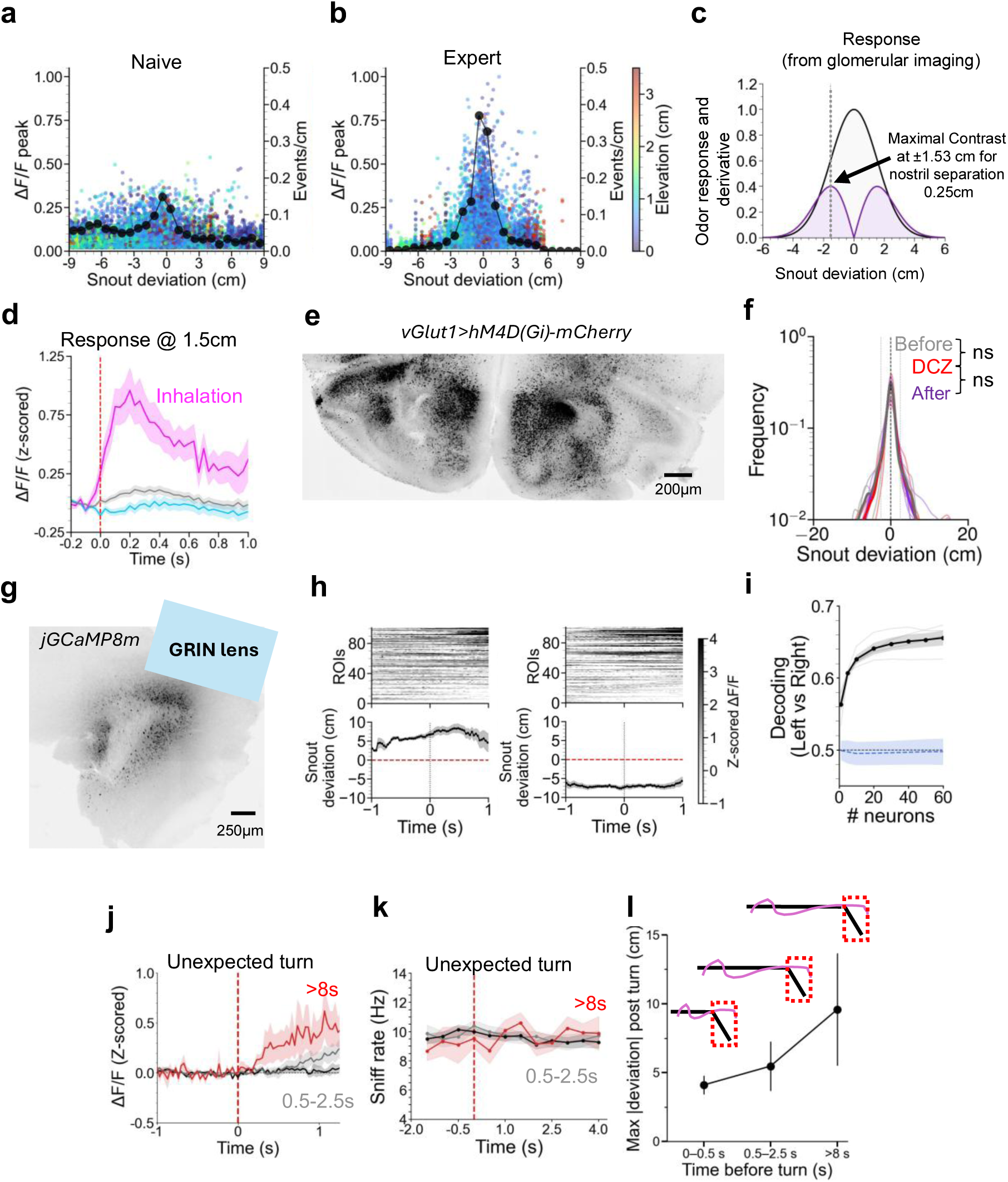
Odor response dynamics during trail tracking. **(a and b)** Glomerular response (ΔF/F peak) and event density as a function of snout deviation in naive (**a**) and expert (**b**) mice. The colormap represents the elevation of the snout. **(c)** Estimated odor concentration profiles modeled as Gaussian-like distributions and mapped through the glomerular response profile from **(b)** to determine the trail width producing maximal bilateral contrast; for a nostril separation of 0.25 cm, the predicted optimum was around 1.5 cm. **(d)** Z-scored responses aligned to inhalation at 1.5 cm. **(e)** Example histology showing bilateral expression of hM4D(Gi)-mCherry in the AON, Scale bar, 200 μm. **(f)** Distributions of snout deviations from 4 mice from the session before (grey), during (DCZ, red) and after (purple) expressing viral vector containing only the fluorophore mCherry (ns, not significant, Mann-Whitney U test). **(g)** Example post-hoc histology of a coronal section showing the lens position in the AON, Scale bar, 250 μm. **(h)** Population activity from the neurons in the AON (top) and corresponding deviation of the snout from the trail (bottom) aligned to time points when the animal was away from the trail (positioned more than 5 cm to either side) from 1 session. **(i)** Decoding probability of the position of the snout relative to the trail as a function of number of neurons from the AON. **(j)** ΔF/F responses from the AON aligned to unexpected bends for short and long pre-turn intervals. **(k)** Sniff rate aligned to the unexpected encounter of a bend in the trail as a function of time following the trail viz. inhalation for short (0.5 s, gray; 0.5–2.5 s, black) and long (>8 s, red) pre-turn intervals. **(l)** Maximum snout deviation from mice with GRIN lenses targeting the AON after a turn as a function of time before turn (n = 5 mice).

